# In vivo single-cell gene editing using RNA electroporation reveals sequential adaptation of cortical neurons to excitatory-inhibitory imbalance

**DOI:** 10.1101/2025.09.13.672380

**Authors:** Georg Kosche, Alex Fratzl, Martin Munz, Botond Roska

## Abstract

The balance between excitatory and inhibitory neurotransmission is fundamental for normal brain function, yet the adaptation of individual neurons to disrupted excitatory-inhibitory balance is not well understood. We developed highly efficient, in vivo RNA electroporation-based single-cell gene editing to investigate neuronal responses to loss of fast inhibition. Using CRISPR-Cas9 components delivered as RNA, we knocked out GABA-A receptor β subunits in individual layer 2/3 cortical neurons in mouse visual cortex, eliminating fast inhibition. In vivo patch-clamp recordings revealed that cortical neurons adapted to inhibition loss through two sequential mechanisms: a transient reduction of excitatory synaptic input, followed by intrinsic membrane property changes that decreased input resistance. This sequential adaptation program ultimately prevented target neurons from contributing spikes to the cortical network. Our RNA-based single-cell gene editing approach enables investigation of cellular responses independent of network effects, providing new insights into neuronal homeostasis and gene function in individual cells in vivo.

## Introduction

Normal brain function depends on maintaining equilibrium between excitatory and inhibitory neurotransmission ((“E/I balance”). Neurons continuously receive and integrate excitatory and inhibitory inputs, and their timing and balance^1–3^ determine whether action potentials are generated and propagated, ultimately driving behavior. E/I balance is dynamically regulated through multiple mechanisms, including neuronal plasticity^4,5^, homeostatic plasticity^6,7^, and metaplasticity^8^, which modulate receptor expression, changes in ion channel function, and neurotransmitter release. This dynamic regulation allows for both stability and flexibility in information processing by neural circuits. A disruption of the brain’s E/I balance can result in neurological diseases such as epilepsy^9,10^ or autism^11,12^.

Our current understanding of how neurons balance excitation and inhibition^3,13–17^ is mostly based on studies involving perturbations that affect extensive brain networks or many cells of a given cell type. When the E/I balance of a large neuronal population is perturbed, two processes can occur simultaneously. First, the E/I input imbalance in individual neurons may be adapted by mechanisms such as homeostatic^6,7,18^ plasticity. Second, the perturbed neuronal network may respond with coordinated activity, such as developing rhythmic activity, which can itself induce further adaptations in participating neurons^19^. Because these processes occur concurrently, the identity and sequence of cellular mechanisms of adaptation to the disruption of E/I balance cannot be fully understood through network-level perturbations.

In vivo single-cell gene editing could provide an approach to study cellular adaptation without network effects by removing inhibitory receptors from individual neurons, followed by in vivo single-cell electrophysiology to measure the consequences of eliminating inhibition. In vivo single-cell electroporation of DNA has enabled labeling and connectivity tracing of individual cortical neurons^20,21^, however, targeted editing of endogenous genes using this approach has been limited to *EGFP*^22^. This limitation likely stems from the variable success rate of DNA electroporation (30-75%^22–24^). During electroporation, DNA must traverse three lipid bilayer membranes — the cell membrane and the double nuclear membrane — to reach the nucleus where transcription occurs. These multiple barriers are likely to contribute to the variable success rates observed with DNA electroporation. In contrast, mRNA only needs to cross the cell membrane to reach the cytoplasm, where translation occurs. In vivo single-cell RNA electroporation for gene expression or editing has not been previously reported, despite recent advances enabling production of highly stable mRNA molecules in vitro^25^. We hypothesized that single-cell mRNA electroporation could provide genetic access to cells in vivo with reliably high success rates, potentially enabling high-efficacy genome editing.

Here, we developed highly efficient in vivo single-cell RNA electroporation and gene editing to investigate how targeted and rapid elimination of GABA-A receptors affects spiking behavior as well as synaptic and intrinsic excitability of mouse cortical neurons. First, we demonstrated that targeted in vivo single-cell electroporation of mRNA encoding tdTomato, EGFP, or Cre recombinase led to corresponding protein expression in selected neurons in mouse cortex in vivo with more than 85% efficacy. In the case of successful electroporation, co-expression of a second gene was 100% efficient. We then showed that when electroporation was successful, evidenced by fluorescent protein expression (>85%), a nearly 100% efficient EGFP knockout in layer 1 (L1) GABAergic cortical neurons was achieved using Cas9 mRNA and two guide RNAs (gRNAs). Next, we electroporated seven RNAs—Cas9, tdTomato, and Cre mRNAs and four gRNAs—to knock down the three β subunits of GABA-A receptors in layer 2/3 (L2/3) neurons in the visual cortex. Using in vivo patch-clamp recordings, we demonstrated that inhibitory input to Cas9-gRNA-electroporated neurons was reduced by 98% within two days. The magnitude of excitatory input to neurons lacking inhibitory input was significantly reduced, also within two days. Despite the marked reduction in excitation, in the absence of inhibition, the spiking activity of the neurons increased in awake animals. Remarkably, the magnitude of the excitatory input returned to its original baseline level one week after electroporation; however, neurons became silent: their spiking activity was significantly reduced. This reduced spiking was due to intrinsic changes in excitability, such as a reduction in membrane input resistance and an increase in the current threshold required to trigger action potentials. Therefore, in contrast to highly increased spiking activity observed after the loss of GABA-A receptor-mediated inhibition within a large cortical network^13,26,27^, we found that single cortical neurons that lose GABA-A inhibition become silent. This is the result of two sequential phases of adaptation: a transient reduction of excitatory synaptic input, followed by a longer lasting intrinsic membrane adaptation. Therefore, the single-cell RNA-based gene editing approach we developed here isolates cellular responses from network confounds, uncovering novel principles of neuronal adaptation and gene function in the intact brain.

## Results

### In vivo single-cell RNA electroporation allows fast and reliable expression of multiple proteins

We first tested single-cell electroporation of mRNA encoding the red fluorescent protein tdTomato into cortical neurons in vivo (Fig. 1A). Thy1-GCaMP6s^28^ mice were anesthetized and head-fixed under a two-photon microscope after cortical window implantation over the left primary visual cortex. Thy1-GCaMP6s mice constitutively express the genetically encoded calcium indicator GCaMP6s in most cortical neurons, which allowed us to monitor any electroporation-induced increase in calcium activity. We targeted a GCaMP6s-expressing (green fluorescent) L2/3 cortical neuron using two-photon shadow imaging and approached it with a glass micropipette filled with electroporation solution containing tdTomato mRNA and Alexa 594 red fluorescent dye. We positioned the micropipette on the membrane of the target neuron and electroporated it by delivering a short, negative voltage pulse train (Fig. 1A), using a voltage amplitude that achieved a transmembrane current of less than 1 µA (Fig. S1). We monitored the success of electroporation by observing accumulation of Alexa 594 within the target cell and an increase in GCaMP fluorescence, which was likely due to transient depolarization of the cell membrane. The presence of GCaMP6s in the target neurons was not required for RNA electroporation, but it facilitated the online assessment of electroporation efficiency. Alexa 594 dye promptly filled the targeted cell, making it fluorescent for a few hours (Fig. S1, Video S1). The electroporation of a neuron took less than one minute, and several cells could be electroporated using the same pipette (Fig. 1B). After single-cell electroporation, we sealed the cranial window using a transparent glass cover, which enabled chronic two-photon fluorescence imaging of the targeted neurons over several weeks (Fig. 1B,C).

**Figure 1:**
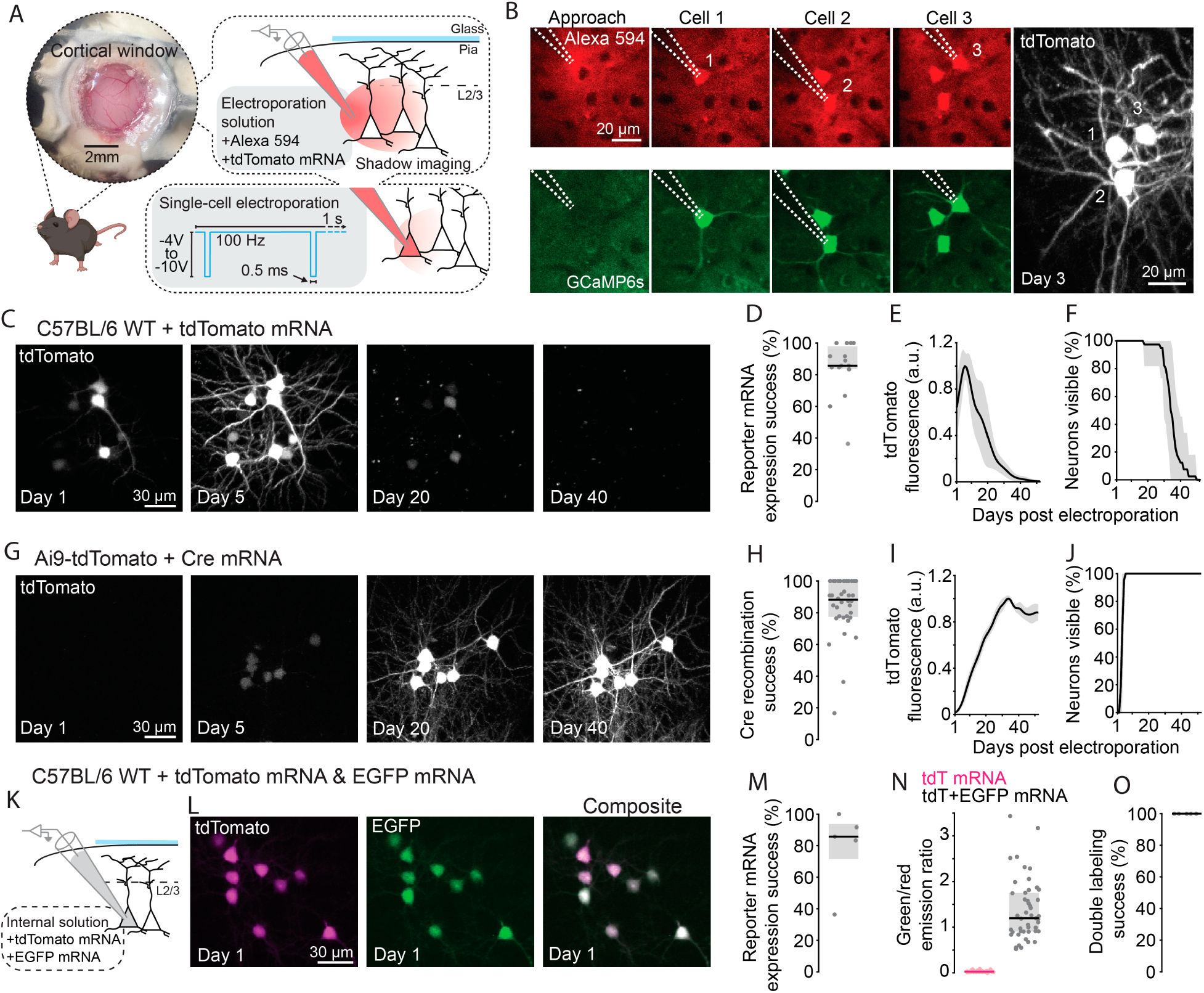
In vivo single-cell RNA electroporation allows fast and reliable expression of multiple proteins A: Schematic of the single-cell RNA electroporation approach. Single-cell RNA electroporation of L2/3 cortical neurons in mice is achieved by targeting cells with a glass micropipette through a cranial window while imaging with a two-photon microscope. **B:** Left panels: Shadow-patch targeting of GCaMP6s^+^ L2/3 neurons in a Thy1-GCaMP6s mouse line during two-photon imaging. Three neurons are shown during the electroporation procedure, which causes transient increases in calcium levels. Alexa 594 is shown in red and GCaMP6s in green. Right panel: Two-photon z-stack max projection showing tdTomato expression in the same three electroporated neurons on the third day after electroporation. **C:** Expression of tdTomato in L2/3 neurons after single-cell electroporation of tdTomato mRNA, imaged over multiple weeks. **D:** Success rate of reporter mRNA electroporation (tdTomato and EGFP combined) quantified on day 1 after electroporation. *n* = 15 mice. Each data point corresponds to an animal. Black like, median; gray shading, interquartile range (IQR) across mice. **E:** Mean tdTomato fluorescence intensity (black line) of electroporated neurons plotted against days since electroporation of reporter mRNA. *n* = 40 neurons from four mice. Gray shading, standard deviation (STD) across neurons. **F:** Mean fraction of visible tdTomato mRNA electroporated neurons (black line), plotted against days since electroporation. Gray shading, STD across neurons. The same cells as in **E**. **G:** Expression of Cre-dependent tdTomato in L2/3 neurons after single-cell electroporation of Cre mRNA, imaged over multiple weeks. **H:** Success rate of Cre mRNA electroporation, quantified by the presence of tdTomato on day 2 after electroporation. *n* = 36 mice. Each data point corresponds to an animal. Black like, median; gray shading, IQR across mice. **I:** Mean tdTomato fluorescence intensity (black line) of electroporated neurons plotted against days since electroporation of Cre mRNA. *n* = 21 neurons from two mice. Gray shading, STD across neurons. **J:** Mean fraction of visible Cre mRNA-electroporated neurons (black line), plotted against days since electroporation. Gray shading, STD across neurons. The same cells as in **I**. **K:** Schematic of the double-labeling using single-cell mRNA electroporation of both EGFP and tdTomato. **L**: Expression of tdTomato (left), EGFP (middle), and both combined (right) in L2/3 neurons after single-cell electroporation of both tdTomato and EGFP mRNA, imaged the day after electroporation. **M**: Success rate of mRNA electroporation quantified by the presence of tdTomato and/or EGPF fluorescence on day 1 after electroporation. *n* = 5 mice. Each data point corresponds to an animal. Black line, median; gray shading, IQR across mice. **N**: Emission ratio between green and red fluorescence in neurons electroporated only with tdTomato mRNA (magenta, *n* = 34 neurons from six mice) versus neurons electroporated with both tdTomato and EGFP mRNA (black, *n* = 47 neurons from five mice). Each data point corresponds to a cell. Magenta and black lines, median; magenta and gray shadings, IQR across neurons. **O**: Double-labeling success rate from the animals displayed in **M**. Each data point corresponds to an animal. Black line, median; gray shading, IQR across neurons.

We observed tdTomato fluorescence (Fig. 1C-F) on the first day after electroporation. Fluorescence peaked at day six (median, interquartile range (IQR): day 5–9, Fig. 1E), after which it decreased with a half-life of 6.11 days (median, IQR: 4.94–7.93 days). We could detect tdTomato-positive neurons for 34.5 days (median, IQR: 32–37 days, Fig. 1F). We achieved similar results when electroporating EGFP mRNA (Fig. S2). EGFP fluorescence peaked on the first day (median, IQR: day 1–1, Fig. S2), had a half-life of 4.00 days (median, IQR: 3.11–4.59 days), and lasted for 21 days (median, IQR: 18–24 days, Fig. S2). The overall success rate of expressing tdTomato or EGFP was 85.71% (median, IQR: 83.65%–97.92%, Fig. 1D). We rarely labeled untargeted neurons or glial cells (two neurons and 12 glial cells in 525 electroporations; Fig. S3). Thus, single-cell electroporation of mRNA encoding fluorescent proteins led to protein expression in more than 85% of electroporated cells, and labeled cells for several weeks.

As a complementary strategy, we tested single-cell RNA electroporation for permanently labeling neurons in the mouse brain. We electroporated mRNA encoding Cre recombinase into cortical neurons of Ai9(RCL-tdT) mice^29^ whose cells express tdTomato only following Cre-mediated recombination. The electroporation of Cre mRNA led to strong expression of tdTomato in target cells (Fig. 1G, Fig. S1), with a high success rate (median: 88.19%, IQR: 77.35%–100%, Fig. 1H) as for mRNAs encoding tdTomato or EGFP. In the case of Cre mRNA electroporation, however, tdTomato fluorescence increased gradually (Fig. 1G,I), with neurons reaching 5% of their maximum fluorescence at day four (median, IQR: day 4–5, Fig. 1I) and peak fluorescence after 34 days (median, IQR: 33–35 days, Fig. 1I). At the end of the chronic imaging experiment (52 days after electroporation), tdTomato fluorescence remained at 86.42% ± 7.05% of peak value (Fig. 1I) and all successfully electroporated cells remained visible (Fig. 1J). Therefore, single-cell electroporation of mRNA encoding Cre recombinase leads to stable, long-lasting labeling of cells that conditionally express a fluorescent protein from their genome.

We next investigated whether single-cell RNA electroporation could lead to the expression of two distinct proteins simultaneously. We performed single-cell electroporation of L2/3 neurons using a micropipette containing two different mRNAs encoding tdTomato or EGFP (Fig. 1K). This led to the expression of both fluorescent proteins as detected one day after electroporation (Fig. 1L). The fluorescent protein expression success rate was as high as in previous experiments (median: 85.71%, IQR: 71.59%–93.75%, Fig. 1M), and every fluorescent neuron expressed both fluorophores simultaneously (Fig. 1N,O). Thus, when two different mRNAs are simultaneously electroporated, the probability of expressing a second protein is 100%, conditional on the expression of the first protein.

### Highly efficient single-cell genomic CRISPR editing in vivo by RNA electroporation

Given the high success rate of expressing proteins using two different RNAs (Fig. 1), we next explored whether single-cell RNA electroporation could be used for genome editing by the CRISPR-Cas9 system, which requires a minimum of two RNAs: Cas9 mRNA and a gRNA. We first aimed to edit—and thereby knock out—the gene driving the endogenous expression of EGFP in a heterozygous mouse line that constitutively expresses EGFP in cortical inhibitory neurons (Gad67-*EGFP*^30^). Heterozygous Gad67-*EGFP* mice have a single copy of the *EGFP* gene in their genome, which allows quantifying single CRISPR/Cas9 editing events based on the disappearance of EGFP fluorescence. We electroporated cortical L1 EGFP-positive neurons in vivo with a micropipette filled with a mixture of gRNA targeting the *EGFP* gene, Cas9 mRNA, and tdTomato mRNA (Fig. 2A). We then tracked electroporated EGFP-positive neurons that also became tdTomato-positive after electroporation (Fig. 2A,B). Within 30 days, 75% of the EGFP-tdTomato double-positive cells (median, IQR: 57.59%–83.93%) lost their EGFP signal (Fig. 2B-D, Fig. S4). In contrast, none of the non-electroporated control cells lost the EGFP signal in the same timeframe (Fig. S4). To control for the potential death of neurons that lost EGFP fluorescence after electroporation, we quantified tdTomato fluorescence in electroporated neurons at different time points following electroporation. There was no difference in tdTomato fluorescence decay between neurons that lost and those that retained EGFP fluorescence (Fig. S4). Thus, the loss of the EGFP signal results from CRISPR editing rather than cell death.

**Figure 2:**
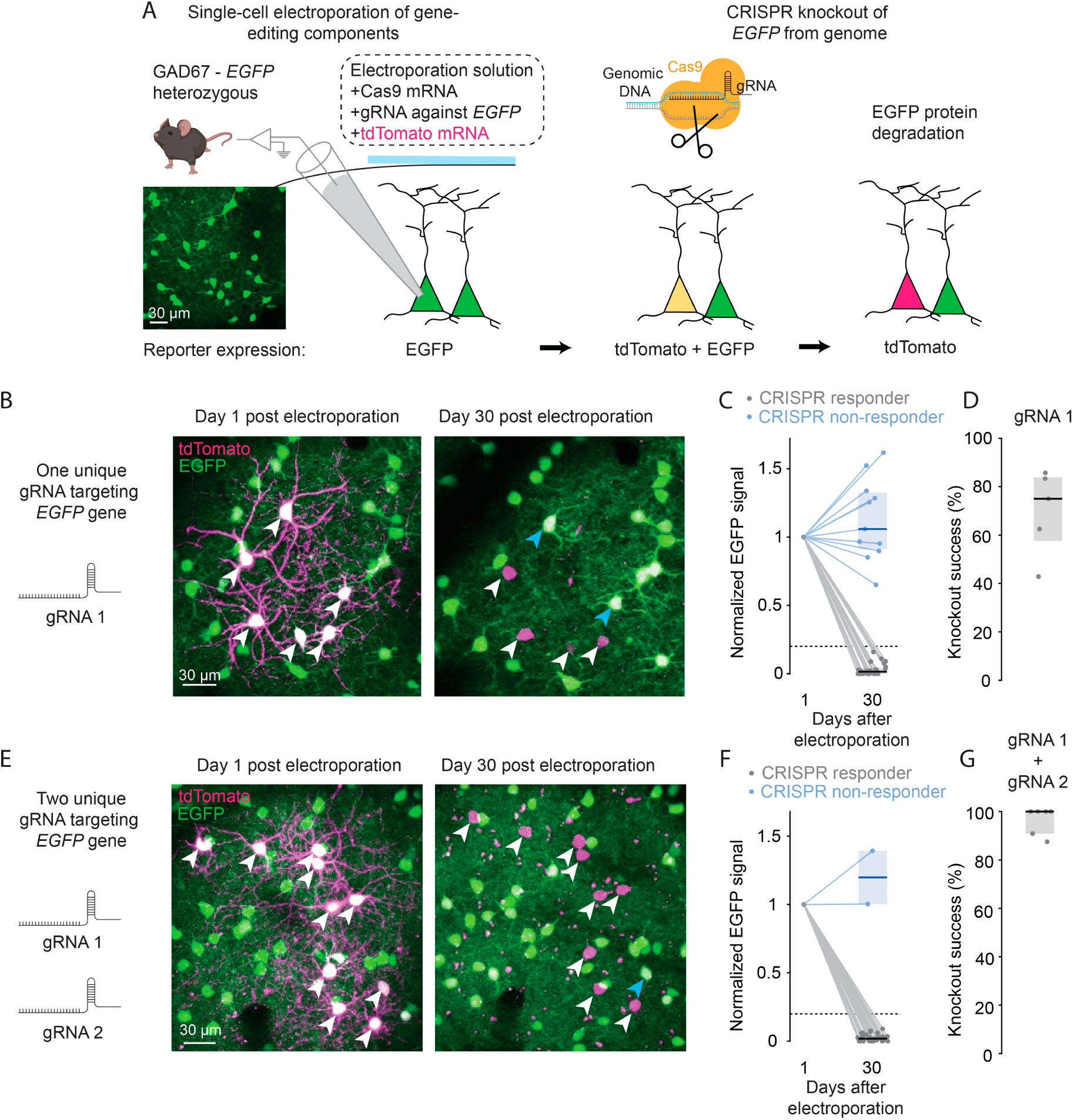
Highly efficient single-cell genomic CRISPR editing in vivo by RNA electroporation A: Schematic of the single-cell genomic CRISPR editing approach using RNA electroporation. tdTomato reporter mRNA, Cas9 mRNA and gRNA against *EGFP* are electroporated in L1 EGFP^+^ GAD67-neurons. Successfully electroporated neurons will express the tdTomato reporter and be double labeled after electroporation. Successful CRISPR editing will cause a progressive loss of EGFP fluorescence over time, due to protein degradation. **B:** Expression of tdTomato and EGFP in EGFP^+^ L1 cortical neurons on day 1 (left) and day 30 (right) after single-cell electroporation of one unique gRNA against EGFP. White arrows demarcate electroporated cells in which CRISPR editing was successful (no EGFP fluorescence remains at day 30); blue arrows demarcate electroporated cells where CRISPR editing was not successful (EGFP fluorescence remains at day 30). **C:** EGFP fluorescence intensity in neurons on day 1 and day 30 after single-cell electroporation of one unique gRNA against *EGFP*. Fluorescence of each cell is normalized to its own day 1 fluorescence. CRISPR editing is considered successful if day 30 fluorescence intensity is less than 20% of day 1 fluorescence intensity (thin gray traces), while CRISPR editing is considered unsuccessful otherwise (thin blue traces). *n* = 36 cells from five imaging sites in four mice. Each data point corresponds to a cell. Black and blue solid lines, median; gray and blue shadings, IQR across cells. **D:** CRISPR editing success rate in electroporated neurons from **C**. *n* = 5 imaging sites in four mice. Each data point corresponds to an imaging site. Black line, median; gray shading, IQR across imaging sites. **E:** Same as **B** but using two different gRNA against *EGFP*. **F:** Same as **C** but using two different gRNA against *EGFP*. *n* = 34 cells from six imaging sites in three mice. **G:** Same as **D** but using two different gRNA against *EGFP*. *n* = 6 imaging sites in three mice.

Since CRISPR/Cas9 editing is an independent event for each editing site, we reasoned that using two distinct gRNAs targeting different locations within the same gene would further increase editing efficiency. To test this, we electroporated EGFP-expressing L1 neurons in heterozygous Gad67-*EGFP* mice with Cas9 mRNA, tdTomato mRNA, and two different gRNAs targeting the *EGFP* gene at different sites (Fig. 2E). Remarkably, this led to the disappearance of EGFP (Fig. 2F) in almost all successfully electroporated (>85%) neurons (median: 100%, IQR: 90.91%– 100%, Fig. 2G), indicating a near-complete genomic *EGFP* knockout rate. Taken together, these results demonstrate that single-cell RNA electroporation using CRISPR-Cas9 components with two different gRNAs can knock out one allele of a gene in cortical neurons in vivo with close to 100% efficacy, provided that electroporation is successful (>85%).

### Single-cell knockout of GABA-A-R β subunits results in rapid loss of fast synaptic inhibition in vivo

We then aimed to use in vivo single-cell RNA electroporation of CRISPR-Cas9 components to knock out endogenous genes that are necessary for forming receptors receiving inhibitory synaptic input in single cortical neurons. We chose the GABA-A receptor, a GABA-gated chloride channel, as GABA is the primary inhibitory neurotransmitter within the neocortex, and GABA-A receptors are responsible for fast synaptic inhibition^31^.

The GABA-A receptor consists of a pentamer of transmembrane subunits of α, β, and γ types^32^ (Fig. 3A). GABA-A receptors are only functional in the presence of β subunits^33–35^. Therefore, we reasoned that knocking out all three genes coding for the β subunits (*GABRB1*, *GABRB2*, and *GABRB3*) would result in complete loss of GABA-A-mediated inhibition^35^. Given the near 100% gene-editing success rates achieved with RNA electroporation using two distinct gRNAs against the same gene (Fig. 2), we targeted each of the three genes with two distinct gRNAs (Fig. 3B, GABA-A CRISPR). We used already characterized gRNAs for each of the three β subunits^36^, and designed an additional gRNA to target a region conserved across all three β subunits. Thus, we electroporated four gRNAs.

**Figure 3:**
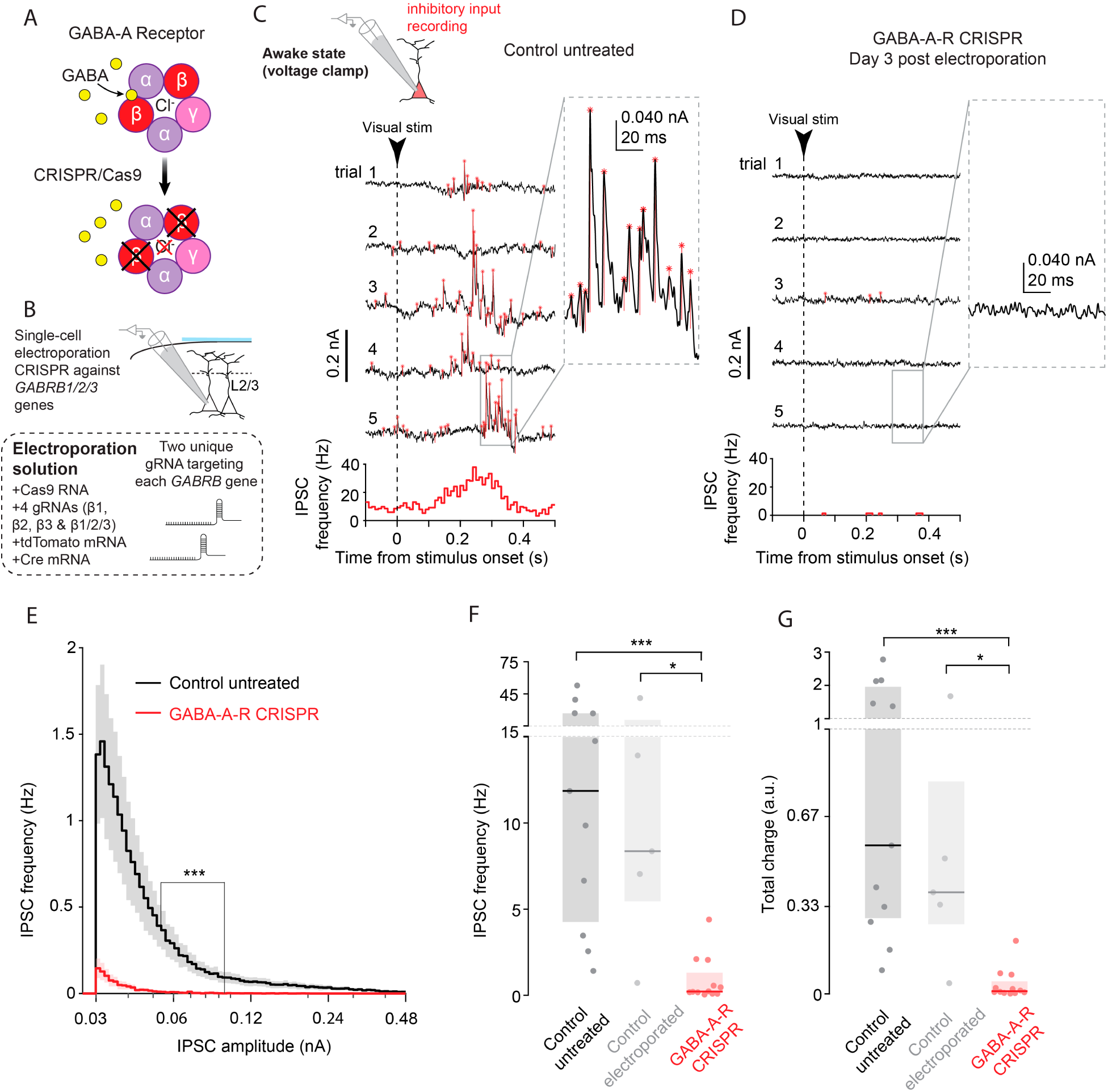
Single cell knockout of GABA-A-R β subunits results in rapid loss of fast synaptic inhibition in vivo A: Schematic of the GABA-A receptor. Beta subunits (shown in red) are targeted by CRISPR/Cas9. Loss of beta subunits prevents assembly of a functional channel, leading to loss of GABAergic chloride (Cl^-^) conductance. **B:** Schematic of the single-cell RNA CRISPR editing approach against the *GABRB1/2/3* genes in L2/3 neurons of Ai9-tdTomato mice. **C:** Awake whole-cell voltage clamp recording of inhibitory post-synaptic currents (IPSCs) of an untreated L2/3 neuron during wakefulness. Shown are five trials of visual stimulus presentation, aligned on stimulus onset, as well as the IPSC peristimulus time histogram (PSTH) for all stimulus presentations (*n* = 153 trials). The inset shows a detailed view of IPSCs (red star = detected event peak, thin red line = event amplitude). **D:** Voltage clamp recording of a GABA-A-R CRISPR electroporated L2/3 neuron (three days after electroporation) during wakefulness. Same as in **C**, PSTH of all detected IPSC events are shown in red (*n* = 68 trials). **E:** Mean population distributions of IPSC strength in control untreated neurons (black, *n* = 11 neurons in seven mice) and GABA-A-R CRISPR electroporated neurons (red, *n* = 12 neurons in seven mice). Gray and red shadings, standard error of the mean (SEM) across neurons. *p* = 5.40 × 10^-19^, two-sample Kolmogorov-Smirnov test. **F:** ISPC frequency of control untreated neurons (black, *n* = 11 neurons in seven mice), control electroporated neurons (gray, *n* = 5 neurons in three mice), and GABA-A-R CRISPR neurons (red, *n* = 12 neurons in seven mice). Each data point corresponds to a cell. Black, gray and red lines, median; gray, light gray and red shadings, IQR across neurons. Control untreated - Control electroporated: *p* = 0.991, Control untreated – GABA-A-R CRISPR: *p* = 3.64 × 10^−4^, Control electroporated - GABA-A-R CRISPR: *p* = 1.79 × 10^−2^, Dunn’s multiple comparison test, preceded by a Kruskal-Wallis test, *p* = 2.30 × 10^−4^. **G**: IPSC charge of control untreated neurons (black, *n* = 11 neurons in seven mice), control electroporated neurons (gray, *n* = 5 neurons in three mice), and GABA-A-R CRISPR neurons (red, *n* = 12 neurons in seven mice). Each data point corresponds to a cell. Black, gray and red lines, median; gray, light gray and red shadings, IQR across neurons. Control untreated - Control electroporated: *p* = 0.937, Control untreated – GABA-A-R CRISPR: p = 1.38 × 10^−4^, Control electroporated - GABA-A-R CRISPR: *p* = 2.27 × 10^−2^, Dunn’s multiple comparison test, preceded by a Kruskal-Wallis test, *p* = 1.16 × 10^−4^.

We aimed to knock out the three β subunits in L2/3 neurons of the visual cortex, as neural activity in these cortical layers is under tight control of local inhibitory interneurons^37,38^. We electroporated neurons in vivo in Ai9(RCL-tdT) mice with a micropipette containing seven RNAs: the four distinct gRNAs together with Cas9 mRNA, tdTomato mRNA, and Cre mRNA (Fig. 3B). We used both tdTomato and Cre mRNA to achieve fast and long-lasting tdTomato labeling (*n* = 32 mice, median success rate: 87.5%, IQR: 77.35%–100%), as we have shown above (Fig. 1). To functionally evaluate the success of knocking out the GABA-A receptors from the target cell, we performed in vivo whole-cell voltage-clamp recordings in awake, head-fixed mice^39^ during ongoing visual stimulation to increase activity. We recorded inhibitory postsynaptic currents (IPSCs) from three groups of neurons (Fig. 3C-G, Fig. S5). The first group consisted of neurons electroporated with all seven RNAs (“GABA-A-R CRISPR electroporated neurons”), the second included neurons electroporated with three RNAs—tdTomato mRNA, Cre mRNA, and Cas9 mRNA, but without gRNAs (“control electroporated neurons”), and the third comprised non-electroporated neurons (“control untreated neurons”).

Remarkably, GABA-A-R CRISPR electroporated neurons expressing tdTomato showed a near-complete absence of IPSCs when recorded two or more days after electroporation (Fig. 3D-G). Control untreated and control electroporated neurons showed IPSC frequencies of 11.86 Hz (median, IQR: 4.26 Hz – 26.80 Hz) and 8.36 Hz (median, IQR: 5.46 Hz – 20.71 Hz), respectively. GABA-A-R CRISPR electroporated neurons showed only a few events that were mostly indiscernible from noise (Fig. 3D-F, Fig. S5; median frequency: 0.21 Hz, IQR: 0.13 Hz – 1.31 Hz), corresponding to a 98% reduction in IPSC frequency compared to control untreated neurons, and more than 97% compared to control electroporated neurons. Similarly, the total inhibitory charge, measured as the product of IPSC frequency and amplitude, was reduced by more than 98% in GABA-A-R CRISPR electroporated neurons compared to control untreated neurons, and more than 97% compared to control electroporated neurons (Fig. 3E,G, Fig. S5). Note, that the recorded decrease in IPSC frequency is likely due to the major decrease in the amplitude of the individual IPSC events (measured as charge), such that the inhibitory input became indistinguishable from noise.

Interestingly, on the first day after electroporation, GABA-A-R CRISPR electroporated neurons still retained control levels of inhibition (Fig. S5), suggesting that the reduction of functional GABA-A channels occurs between day one and two after electroporation. Taken together, single-cell RNA electroporation of Cas9 mRNA and the four gRNAs targeting the three GABA-A β subunits eliminated fast synaptic inhibitory input to L2/3 cortical neurons in vivo.

### The immediate effect of loss of fast synaptic inhibition is increased spiking activity

The rapid elimination of inhibition from GABA-A-R CRISPR electroporated neurons enabled us to investigate how individual neurons adapt to this perturbation of the E/I balance. Since fast inhibition was abolished by day 2 after electroporation (Fig. 3, Fig. S5), we first asked how the spiking activity changed at days 2-3. We measured the membrane potential of individual L2/3 neurons in the primary visual cortex using the current-clamp mode of whole-cell patch clamp in awake mice, while presenting visual stimuli to increase activity. Here we compared two groups of neurons: control untreated neurons and GABA-A-R CRISPR electroporated neurons two-three days after electroporation.

In awake mice, control untreated neurons underwent transient, network-driven, membrane potential fluctuations, alternating between hyperpolarized and depolarized states, and generated action potentials during the depolarized states as described before^40,41^ (Fig. 4A). Consistent with reports of network effects of removing inhibition^10,13^, GABA-A-R CRISPR electroporated neurons recorded two-three days after electroporation had significantly higher spiking activity than control untreated neurons (Fig. 4A-B; control untreated neurons, median: 0.27 Hz, IQR: 0.03 Hz–0.36 Hz; GABA-A-R CRISPR electroporated neurons, median: 1.21 Hz, IQR: 0.56 Hz – 1.61 Hz).

**Figure 4:**
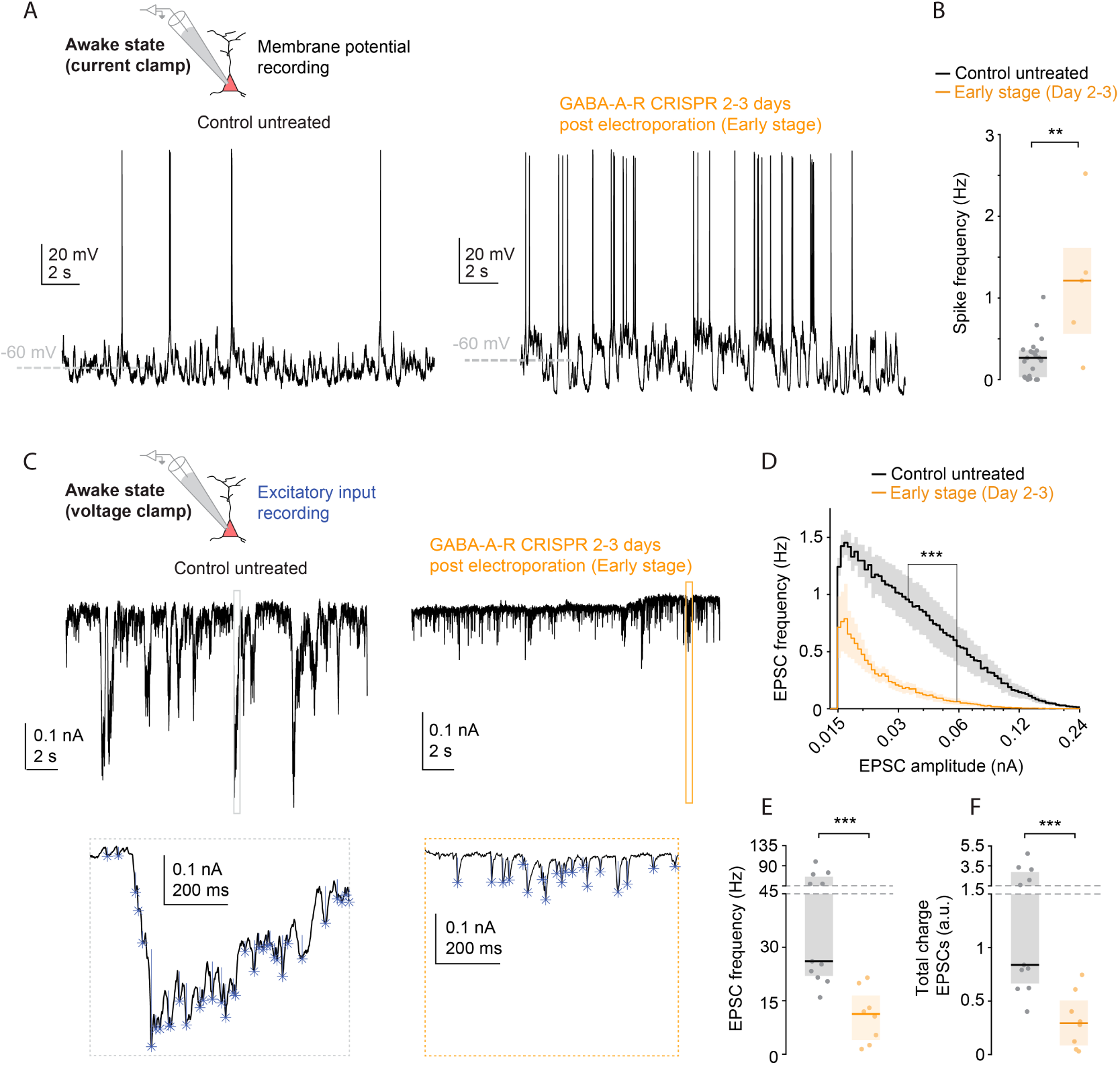
The immediate effect of loss of inhibition is increased spiking activity and decreased excitatory input A: Top: Awake whole-cell current clamp recording of the membrane potential of a control untreated (left) and a GABA-A-R CRISPR (right, two-three days after electroporation, early-stage) L2/3 neuron. **B:** Spike rate of untreated neurons (black, *n* = 21 neurons in nine mice) and early-stage GABA-A-R CRISPR neurons (yellow, *n* = 5 neurons in two mice). Each data point corresponds to a cell. Black and yellow lines, median; gray and yellow shadings, IQR across neurons. *p* = 9.67 × 10^−3^, Wilcoxon rank-sum test. **C:** Top: Awake whole-cell voltage clamp recording of excitatory post-synaptic currents (EPSCs) of a control untreated (left) and an early-stage GABA-A-R CRISPR (right) L2/3 neuron during wakefulness. Bottom: insets showing a detailed view of EPSCs (blue star = detected event peak, thin blue line = event amplitude). **D:** Mean population distributions of EPSC amplitude in control untreated neurons (black, *n* = 11 neurons in seven mice) and early-stage GABA-A-R CRISPR electroporated neurons (yellow, *n* = 8 neurons in four mice). Gray and yellow shadings, SEM across neurons: *p* = 6.05 × 10^-8^, two-sample Kolmogorov-Smirnov test. **E:** EPSC frequency of untreated neurons (black, *n* = 11 neurons in seven mice) and early-stage GABA-A R CRISPR neurons (yellow, *n* = 8 neurons in four mice). Black and yellow lines, median; gray and yellow shadings, IQR across neurons. *p* = 1.85 × 10^-4^, Wilcoxon rank-sum test. **F**: EPSC charge of untreated neurons (black, *n* = 11 neurons in seven mice) and late-stage GABA-A-R CRISPR neurons (yellow, *n* = 8 neurons in four mice) neurons. Each data point corresponds to a cell. Black and yellow lines, median; gray and yellow shadings, IQR across neurons. *p* = 5.03 × 10^-4^, Wilcoxon rank-sum test.

### Early adaptation to loss of inhibition downregulates excitatory input

To understand whether there was an adaptation mechanism that limited the increase in spiking activity, we recorded excitatory postsynaptic currents (EPSCs) in individual L2/3 cortical neurons using the voltage-clamp mode of whole-cell patch clamp in awake animals two-three days after electroporation. Voltage clamp recording of GABA-A-R CRISPR electroporated neurons at this stage showed significantly reduced excitatory synaptic input (Fig. 4C-F). Both EPSC frequency (control untreated neurons, median: 26.07 Hz, IQR: 21.97 Hz – 65.51 Hz; GABA-A-R CRISPR electroporated neurons, median: 11.30 Hz, IQR: 3.98 Hz – 16.49 Hz, Fig. 4E) and total excitatory charge (control untreated neurons, median: 0.84 a.u., IQR: 0.66 a.u. – 2.88 a.u.; GABA-A-R CRISPR electroporated neurons, median: 0.29 a.u., IQR: 0.09 a.u. – 0.51 a.u., Fig. 4F) were approximately half compared to control untreated neurons. This suggests that at the time when inhibition was rapidly decreasing, excitatory input in L2/3 cortical neurons was rapidly downregulated. However, this rapid decrease in excitation did not prevent the increase of firing in neurons lacking fast inhibition (Fig. 4A).

### The late result of loss of inhibition is neuronal silencing

It is possible that excitatory input of a neuron that loses inhibition is downregulated, but this process is not complete within two-three days and therefore cannot prevent an increase in spiking. Therefore, we measured the membrane potential of GABA-A-R CRISPR electroporated neurons in awake mice seven or more days after electroporation. Indeed, by this time, GABA-A-R CRISPR electroporated neurons were almost completely silent, producing only few action potentials (Fig. 5A-B, control untreated neurons, median: 0.27 Hz, IQR: 0.03 Hz – 0.36 Hz; GABA-A-R CRISPR electroporated neurons, median: 0 Hz, IQR: 0 Hz – 0.06 Hz).

**Figure 5:**
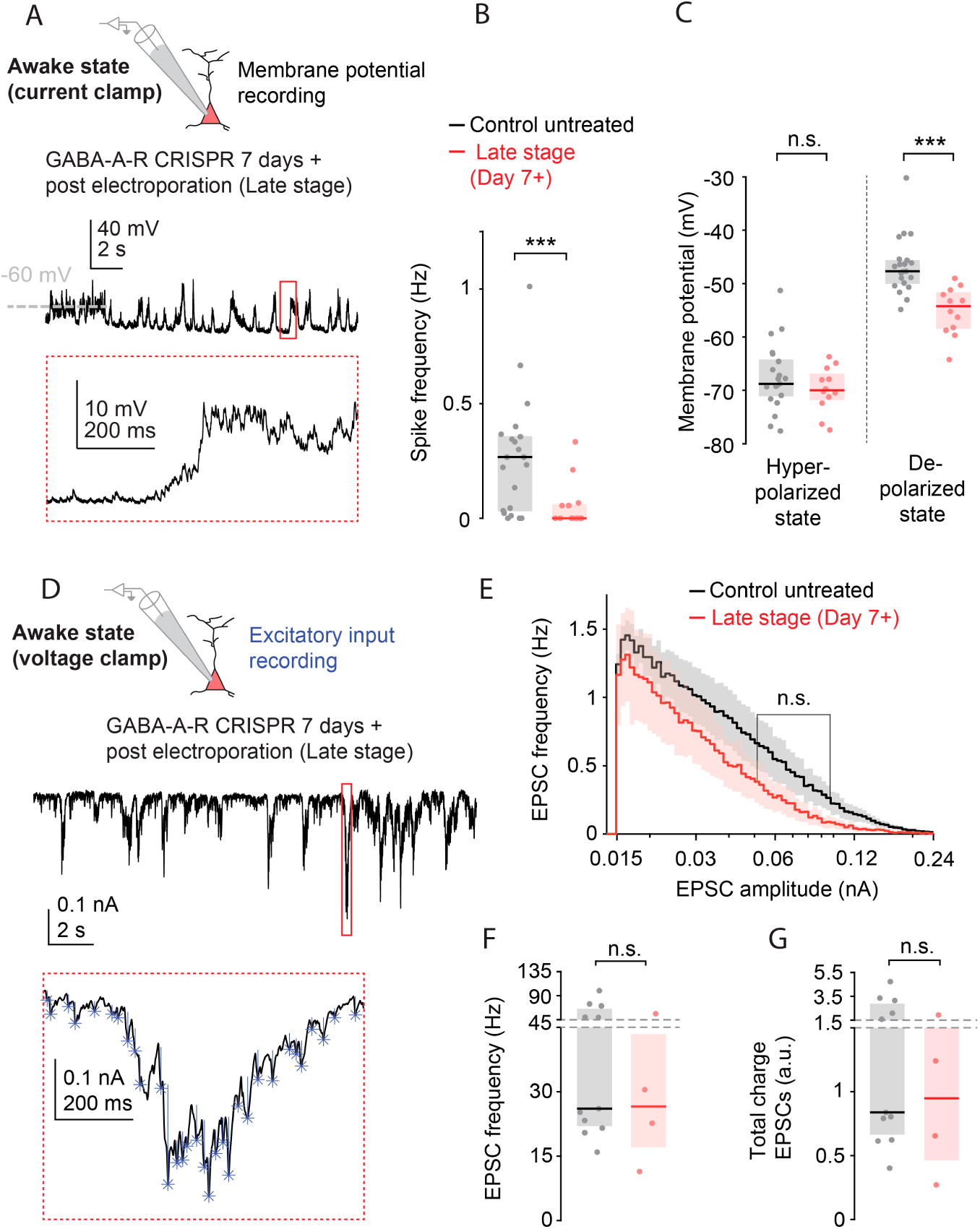
The late result of loss of inhibition is neuronal silencing without a change in excitatory input A: Top: Awake whole-cell current clamp recording of the membrane potential of a late-stage GABA-A-R CRISPR (more than seven days after electroporation) L2/3 neuron. Bottom: inset showing a detailed view of the membrane potential fluctuations. **B:** Spike rate of untreated neurons (black, *n* = 21 neurons in nine mice) and late-stage GABA-A-R CRISPR neurons (red, *n* = 12 neurons in five mice). Each data point corresponds to a cell. Black and red lines, median; gray and red shadings. IQR across neurons. *p* = 4.42 × 10^−3^, Wilcoxon rank-sum test. **C**: Hyperpolarized (left) and depolarized (right) membrane potentials of untreated neurons (black, *n* = 21 neurons in nine mice) and late-stage GABA-A-R CRISPR neurons (red, *n* = 12 neurons in five mice). Each data point corresponds to a cell. Black and red lines, median; gray and red shadings. IQR across neurons. Hyperpolarized state: *p* = 0.291, depolarized state: *p* = 8.75 × 10^−6^, Wilcoxon rank-sum test. **D:** Top: Awake whole-cell voltage clamp recording of EPSCs of a late-stage GABA-A-R CRISPR L2/3 neuron (more than seven days after electroporation). Bottom: inset showing a detailed view of EPSCs (blue star = detected event peak, thin blue line = event amplitude). **E:** Mean population distributions of EPSC amplitude in control untreated neurons (black, *n* = 11 neurons in seven mice) and late-stage GABA-A-R CRISPR electroporated neurons (red, *n* = 4 neurons in three mice). Gray and red shadings, SEM across neurons. *p* = 0.125, two-sample Kolmogorov-Smirnov test. **F:** EPSC frequency of untreated neurons (black, *n* = 11 neurons in seven mice) and late-stage GABA-A-R CRISPR neurons (red, *n* = 4 neurons in three mice). Each data point corresponds to a cell. Black and red lines, median; gray and red shadings, IQR across neurons. *p* = 0.571, Wilcoxon rank-sum test. **G:** EPSC charge of untreated neurons (black, *n* = 11 neurons in seven mice) and late-stage GABA-A-R CRISPR neurons (red, *n* = 4 neurons in three mice). Each data point corresponds to a cell. Black and red lines, median; gray and red shadings, IQR across neurons. *p* = 0.489, Wilcoxon rank-sum test.

Furthermore, during the natural membrane potential fluctuations, which alternate between hyperpolarized and depolarized states, GABA-A-R CRISPR electroporated neurons and control untreated neurons had hyperpolarized states with similar membrane potentials (Fig. 5C, control untreated neurons, median:-68.78 mV, IQR:-71.11 mV –-64.23 mV; GABA-A-R CRISPR electroporated neurons: median:-69.99 mV, IQR:-71.82 mV –-66.86 mV). However, in their depolarized state, the membrane potential of GABA-A-R CRISPR electroporated neurons was significantly less than that of control untreated neurons (Fig. 5C, control untreated neurons, median:-47.69 mV, IQR:-50.02 mV –-45.59 mV; GABA-A-R CRISPR electroporated neurons: median:-54.24 mV, IQR:-58.49 mV –-51.64 mV). This reduced depolarization is consistent with the loss of firing recorded in GABA-A-R CRISPR electroporated neurons.

### Late adaptation reduces membrane excitability without changing excitatory input

A natural explanation for the recorded loss of spiking activity in GABA-A-R CRISPR electroporated neurons a week after the electroporation is that L2/3 cortical neurons reacted to the increase in spiking at days two-three by an even greater reduction in excitatory inputs. To test this hypothesis, we recorded EPSCs in individual L2/3 cortical neurons in awake animals seven or more days after electroporation.

Remarkably, and in contrast to neurons two-three days after electroporation, GABA-A-R CRISPR electroporated neurons and control untreated neurons now had similar EPSC frequencies (Fig. 5D-F, control untreated neurons, median: 26.07 Hz, IQR: 21.97 Hz – 65.51 Hz; GABA-A-R CRISPR electroporated neurons, median: 26.57 Hz, IQR: 17.04 Hz – 43.40 Hz) and total charges (Fig. 5G, control untreated neurons, median: 0.84 a.u., IQR: 0.66 a.u. – 2.88 a.u.; GABA-A-R CRISPR electroporated neurons, median: 0.95 a.u., IQR: 0.46 a.u. – 1.58 a.u.). Therefore, excitatory input returned to normal levels by one week after electroporation, and a delayed adaptation mechanism was induced that led to reduced firing in L2/3 cortical neurons after the removal of their GABA-A inhibitory input.

A possible delayed adaptation mechanism is a change in intrinsic membrane properties, such that L2/3 cortical neurons respond less to the same excitatory currents. To test this, we recorded the membrane potential of individual L2/3 cortical neurons in the current-clamp mode of whole-cell patch clamp in awake animals seven or more days after electroporation. We assessed the response of the membrane potential to controlled current injections (Fig. 6A).

**Figure 6:**
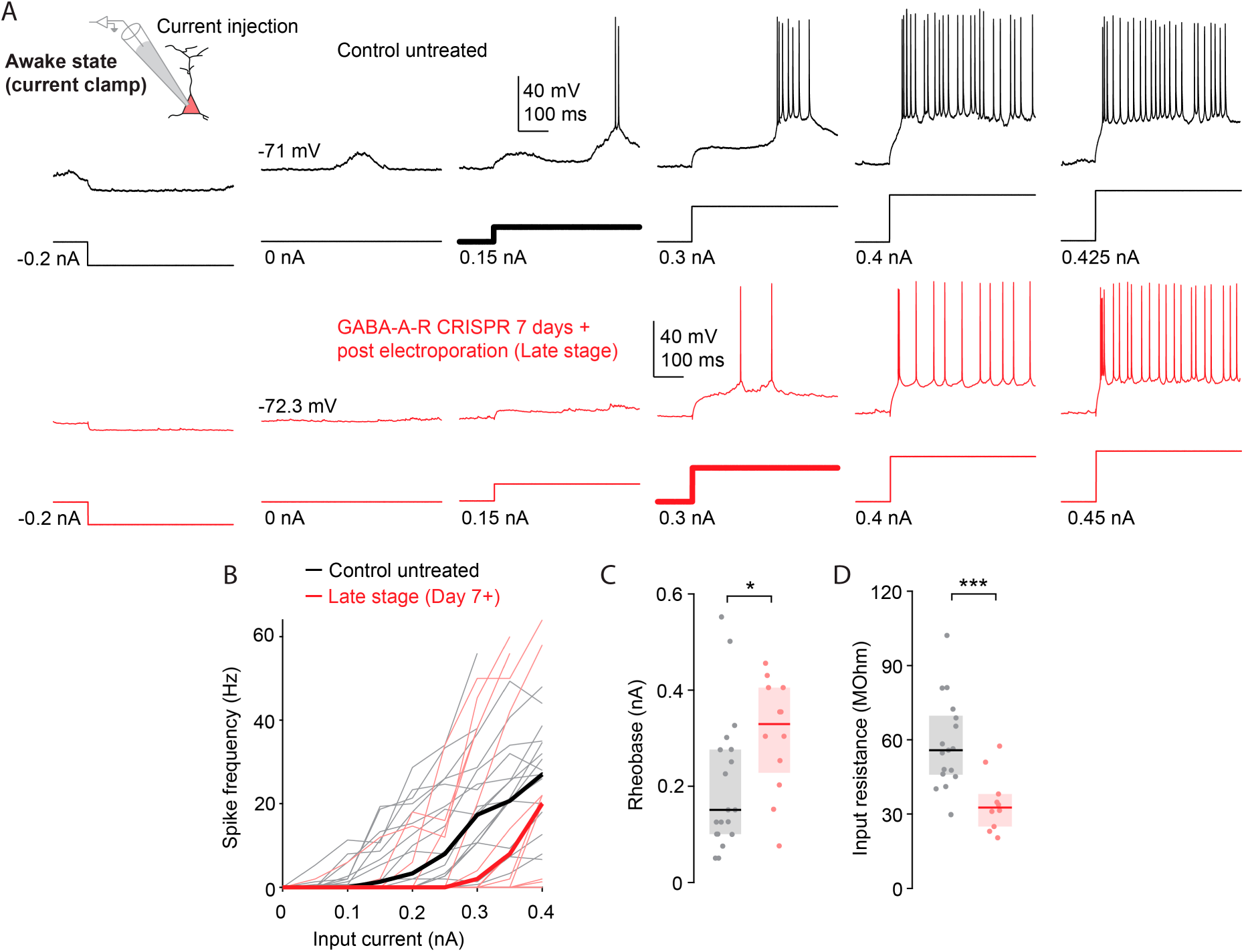
Late adaptation reduces membrane excitability A: Awake whole-cell current clamp recording of the membrane potential of untreated (top row, black) and late-stage GABA-A-R CRISPR L2/3 neurons (more than seven days after electroporation, bottom row, red) during current injections. The corresponding current step trace is displayed below the membrane potential trace. Bold current trace indicates rheobase. **B:** Spike rate of untreated neurons (black, *n* = 19 neurons in nine mice) and late-stage GABA-A-R CRISPR neurons (red, *n* = 12 neurons in five mice) in response to current injections. Bold line, median; each pale line corresponds to a cell. **C:** Rheobase of untreated neurons (black, *n* = 19 neurons in nine mice) and late-stage GABA-A-R CRISPR neurons (red, *n* = 12 neurons in five mice). Each data point corresponds to a cell. Black and red lines, median; gray and red shadings, IQR across neurons. *p* = 1.75 × 10^−2^, Wilcoxon rank-sum test. **D:** Input resistance of untreated neurons (black, *n* = 19 neurons in nine mice) and late-stage GABA-A-R CRISPR neurons (red, *n* = 12 neurons in five mice). Each data point corresponds to a cell. Black and red lines, median; gray and red shadings, IQR across neurons. *p* = 9.16 × 10^−4^, Wilcoxon rank-sum test.

In GABA-A-R CRISPR electroporated neurons, current injections with the same amplitudes resulted in fewer spikes than in control untreated neurons (Fig. 6B). Furthermore, the minimum current needed to elicit a spike (rheobase) was significantly higher in GABA-A-R CRISPR electroporated neurons than in control untreated neurons (Fig. 6C, control untreated neurons, median: 0.15 nA, IQR: 0.1 nA – 0.28 nA; GABA-A-R CRISPR electroporated neurons, median: 0.33 nA, IQR: 0.23 nA – 0.41 nA). Finally, GABA-A-R CRISPR electroporated neurons showed a significantly lower input resistance than control untreated neurons (Fig. 6D, control untreated neurons, median: 55.79 MΩ, IQR: 46.90 MΩ – 69.74 MΩ; GABA-A-R CRISPR electroporated neurons, median: 32.61 MΩ, IQR: 25.01 – 38.10 MΩ). These findings suggest that reduced membrane excitability is the cause of the reduced firing rate in L2/3 cortical neurons seven or more days after the elimination of GABA-A inhibition in vivo.

Notably, the input resistance and rheobase values of GABA-A-R CRISPR electroporated neurons recorded at an earlier stage, i.e., two-three days after electroporation, did not differ from control untreated neurons (Fig. S6, median input resistance: 41.94 MΩ, IQR: 39.63 MΩ – 88.61 MΩ, median rheobase: 0.1 nA, IQR: 0.038 nA MΩ – 0.23 nA). Moreover, at two-three days after electroporation, the membrane potentials of GABA-A-R CRISPR neurons and control untreated neurons did not differ significantly (Fig. S6, GABA-A-R CRISPR electroporated neurons, hyperpolarized state, median:-71.61 mV, IQR:-73.05 mV –-66.38 mV, depolarized state, median:-45.46 mV, IQR:-49.18 mV –-42.13 mV). Therefore, changes in membrane excitability started later than days 2-3 after electroporation.

## Discussion

We have demonstrated that single-cell electroporation using RNA is a highly efficient approach for transient and long-term expression of desired proteins in cortical neurons in vivo. Furthermore, it enables in vivo single-cell gene editing of targeted neurons with high success rates. Electroporation of Cas9 mRNA and gRNAs that target the β subunits of GABA-A receptors in single cortical neurons eliminated fast inhibitory input to selected cortical L2/3 neurons within two days in vivo. This rapid perturbation of E/I balance led to major and unexpected changes in the physiology of L2/3 cortical neurons. Instead of a strong overexcitation that would be expected based on the network effect of removing inhibition^10,13^, individual L2/3 cortical neurons in awake mice became almost silent seven days after electroporation. In effect, GABA-A knockout caused the shut-down of the spiking output of neurons and they became functionally excluded from the network.

Our experiments revealed two sequential adaptive mechanisms. First, a rapid adaptation occurred within two days of electroporation, coinciding with the complete loss of GABA-A inhibition. At this initial phase of adaptation, the excitatory input to individual L2/3 neurons was reduced by half. However, in the absence of fast inhibition, even this reduced excitation led to increased spike rates. Second, a slower adaptation emerged, characterized by changes in intrinsic membrane properties, i.e., specifically decreased input resistance and increased rheobase. By seven days post-electroporation, this second adaptation silenced neuronal spiking despite the fact that excitatory input strength returned to baseline levels. These results demonstrate that loss of fast inhibitory input in individual cortical neurons initiates a sophisticated, sequential adaptation program. Spiking increased in the early phase of adaptation, while membrane potential fluctuations remained unchanged compared to control cells (Fig. S6). This could be explained by a lack of shunting mechanisms of GABAergic inhibition^42,43^, resulting in depolarizations having a higher spike generation potential. The reduction in excitatory input in the early phase likely reflects excitatory synapse plasticity, which has been observed and studied in sensory cortices using deprivation experiments^44,45^. The site of adaptation can be postsynaptic or locally presynaptic in the terminals of axons that provided excitatory input^46^. The output of these terminals may be adjusted as a result of retrograde signaling from the postsynaptic cell^46^. The reduction in excitatory input could be an early mechanism that restores E/I balance within GABA-A receptor depleted neurons. However, when synaptic rebalancing proves insufficient, intrinsic membrane plasticity is initiated. Membrane property adaptations have been documented in response to chronic activity manipulations^15,18^, and appear to be a more definitive solution when synaptic mechanisms fail to normalize neuronal output. It is possible that heightened spiking activity was sensed by the cell which then triggered the intrinsic membrane property changes responsible for neuronal silencing. Observation of these two mechanistically and temporally different adaptation mechanisms required temporally precise perturbation of inhibition in individual neurons, which was enabled by RNA-electroporation-based single-cell gene editing.

Single-cell in vivo delivery of genetic material for applications including cellular labelling^23,47^, optogenetic control^24^, and circuit tracing^20,21^ has traditionally relied on DNA plasmid electroporation. However, that approach has variable success rates of 30-75%^22–24^, with low reliability when multiple plasmids are required. For example, single-cell circuit tracing experiments report success rates of only ∼12% in vitro and 33% in vivo^48^. We demonstrate here that in vivo single-cell electroporation of mRNA achieves substantially higher expression efficiency, exceeding 85%. Critically, co-electroporation of two mRNAs resulted in the same protein expression success rate as a single mRNA delivery, indicating that, conditional on the expression of the first gene, the expression efficiency of other genes is ∼100%. This advance in increasing the efficacy of expression of multiple genes enables reliable implementation of complex methodologies, such as in vivo single-cell genetic engineering, which typically requires efficient co-delivery of multiple guide RNAs alongside protein-coding sequences. Previous attempts at Cas9-mediated single-cell editing in vivo have been limited. Only one study has reported this approach^22^, where DNA plasmid electroporation to knock out EGFP in cortical neurons in mice had a 43% success rate. In contrast, our mRNA-based method achieved EGFP knockdown in ∼100% of successfully electroporated cells (>85% electroporation success rate) using Cas9 mRNA plus two EGFP-targeting gRNAs. We obtained similarly robust results when combining seven RNAs to knock out endogenous GABA-A channels, including four gRNAs targeting β subunits of the GABA-A receptor. Electroporated cortical neurons showed a 98% reduction in IPSC frequency and charge compared to untreated controls.

In vivo single-cell RNA electroporation opens new possibilities for experimentation in neuroscience. First, single-cell mRNA electroporation is fast and the same pipette can be used to target many different cells sequentially for electroporation (Video S1). This approach could enable delivery of genetically encoded activity modulators^49,50^, sensors^51^, or other genes to specific neurons that have been functionally characterized using in vivo two-photon imaging. This would provide genetic control over individual neurons that is unattainable with conventional viral or transgenic approaches. Second, because editor proteins such as Cas9 are rapidly translated from mRNA, the only limiting factor in single-cell gene editing with RNA will be the turnover rate of the target protein within the cell. In our study, we observed reliable editing effects on both reporter proteins and endogenous GABA-A channels within days of electroporation. This contrasts sharply with in vivo gene editing using adeno-associated viral vectors^36^, which requires weeks to achieve editing and provides no control over the precise timing of editing. Recent advances in RNA synthesis technology have made studies using custom mRNA molecules significantly more affordable, and single-cell RNA electroporation enables relatively straightforward experimentation with rapidly evolving gene editing tools^52^. Third, while conventional viral or combinatorial genetic engineering approaches can modify many neurons within a brain region, such methods have an inherent limitation: it becomes difficult to determine whether observed phenotypes arise from intrinsic responses of the edited neurons themselves or from network-level effects caused by simultaneous editing of large numbers of neurons. The reliable single-cell gene editing via RNA electroporation introduced here could provide complementary insights when used alongside conventional editing methods, particularly for understanding neural disorders characterized by network dysfunction, such as autism or epilepsy. Fourth, recent brain-wide single-cell RNA sequencing efforts^53^ have identified numerous cell type-specific genes across thousands of distinct brain cell types. The highly efficient single-cell gene editing approach introduced here provides a method to investigate how these genes contribute to individual cell functions in vivo, offering direct functional validation of transcriptomic findings. Of particular interest are genes associated with neurological or psychiatric diseases.

## Acknowledgments

We thank A. Muller, M. Wang, S. Curtoni and B. György for advice and help with CRISPR gRNA design. We thank C. Pivetta and the IOB animal caretakers. We thank Z. Raicz for visual stimulation software coding and equipment assistance. We thank T. Ayupov, T. Dalmay, M. Moissidis, S. Nadeau, T. Rodrigues and H. Stabb for comments on the manuscript, and S. Nadeau for extended discussions. We thank P. King for English proofreading. This work was supported by a HFSP fellowship (LT000384/2017-L) to G.K., an EMBO fellowship (ALTF 688-2022) to A.F. and Swiss National Science Foundation Synergia grant (CRSII3_141801), European Research Council advanced grant (HURET N°883781), Louis-Jeantet Foundation award, Körber Foundation award, Swiss National Science Foundation grant (31003A_182523), and the NCCR ‘Molecular Systems Engineering’ to BR.

## Author Contributions

**GK**: conceptualization (single-cell gene editing of GABA-A receptor), methodology, formal analysis, investigation (single-cell electroporation, whole-cell current and voltage clamp recordings), data curation, writing original draft, review and editing, visualization. **AF**: methodology, software, formal analysis, investigation (single-cell electroporation), data curation, writing original draft, review and editing, visualization. **MM**: conceptualization (original idea of single-cell electroporation of mRNA), methodology. **BR**: conceptualization, resources, writing original draft, review and editing, supervision.

## Methods

### Animals

Animal experiments were approved by the Veterinary Department of Basel-Stadt and were conducted according to standard ethical guidelines (European Communities Guidelines on the Care and Use of Laboratory Animals, 89/609/EEC). The following transgenic mouse lines of C57BL/6 background were used in this study: Ai9(RCL-tdT)^29^ (JAX stock #007909, The Jackson Laboratory), Ai9(RCL-tdT) crossed with Thy1-GCaMP6s^28^ (JAX stock #024275, The Jackson Laboratory), and Gad67-EGFP^30^ (JAX stock # 007677, The Jackson Laboratory). Male and female adult mice were housed and maintained on a 12-hour light/dark cycle in a pathogen-free animal room, with unlimited access to food and drinking water. Only adult mice were used for experiments.

### Surgical procedures and cortical window implantation

Animals were anaesthetized by intraperitoneal injection of FMM mix (Fentanyl 0.05 mg/kg BW, Medetomidine 0.5 mg/kg BW and Midazolam 5 mg/kg BW). After mice became unresponsive to toe pinch, they were placed on a heat pad (Supertech) and fixed on a surgery apparatus (Narishige). The scalp was carefully cleared of fur using a shaver, local analgesia was administered (Bupivacaine 0.25% + Lidocaine 0.1-0.2%), and corneas were protected with eye gel (Ocutect gel). The scalp was cut above posterior cortex and a small, 1-mm-thick titanium head-holder ring attached using cyanoacrylate glue and dental cement (SuperBond). The skull was then carefully thinned and drilled in a 4-mm-diameter circle above the visual cortex. Care was taken to keep the dura intact while removing the bone, and the exposed cortex was kept moist using cortex buffer containing 125 mM NaCl, 5 mM KCl, 10 mM glucose, 10 mM HEPES, 2 mM MgSO4, and 2 mM CaCl_2_. The cortex was then covered with a 4-mm-diameter glass coverslip (Warner Instruments), which was fixed using cyanoacrylate glue.

### Cortical window access for single cell electroporation and whole-cell recordings

To gain access to layer 2/3 cells with patch-pipettes, mice with cranial windows were anaesthetized briefly using isoflurane (for awake whole-cell recordings), or FMM mix for single-cell electroporation, and the cortical window removed carefully using a dental drill. A 4-mm-diameter coverslip was cut in half using a diamond knife. The area of interest was located under a stereoscope (Leica) and the straight edge of the half-circular window was placed just above it on the cortex, such that access by patch pipette was granted perpendicular to the edge of the glass. The corners of this coverslip were fixed to the skull using biocompatible, UV curable glue (Norland Products), ensuring that the brain underneath was stabilized and not deformed due to movements of the patch pipette. After single-cell electroporation, UV curable glue and the half-coverslip were easily removed with forceps, and a new 4-mm window to allow chronic two-photon imaging was fixed using cyanoacrylate glue to cover the whole craniotomy.

### Single-cell RNA electroporation of L2/3 neurons in mouse visual cortical

#### RNA containing electroporation solution

tdTomato mRNA, *Streptococcus pyogenes* SF370 Cas9 mRNA (Cat. Nr. L-7606), EGFP mRNA (Cat. Nr. L-7601), and Cre mRNA (5moU modified, Cat. Nr. L-7211) were synthesized by Trilink Biotechnologies ordered from Tebubio and delivered in concentrations of 1-2 mg/mL, frozen in 1 mM sodium citrate pH 6.4. All ordered mRNA were capped using Trilink CleanCap, described as a Cap 1 structure. Additionally, all ordered Trilink mRNA was polyadenylated to mimic fully processed mammalian mRNA. Upon receipt, stock solutions were thawed and aliquoted to one-use 3-μL aliquots, refrozen and stored at-80 C until the day of the experiment. Guide RNAs (Integrated DNA Technologies) for CRISPR/Cas9 experiments were ordered as sgRNA and similarly aliquoted and stored.

#### gRNA target sequences

GABRB1 forward (5′ to 3′) GCCGCGAGGGCTTCGGGCGT; GABRB2 forward (5′ to 3′) CAGACAGCGGCGATTATTAA; GABRB3 forward (5′ to 3′): ACGGTCGACAAGCTGTTGAA. Additionally, a guide RNA was designed to target a conserved region of the three GABA-A receptor beta subunits: GABRB Conserved forward (5′ to 3′): CGGATGACATCGAATTTTAC. EGFP target sequence 1 forward (5′ to 3′): GGCGAGGGCGATGCCACCTA. EGFP target sequence 2 forward (5′ to 3′): GCTGAAGCACTGCACGCCGT. TdTomato target sequence forward (5′ to 3′): TACAGGGTCCGCTTCCCTG.

On the day of the experiment, electroporation solution was prepared by mixing 0.4 M K-Gluconate (Sigma), RNAse-free water (Sigma), 25 µM Alexa Fluor 594 Hydrazide, and the required RNA aliquots, followed by filtration through a 0.45-µm centrifugation filter (Costar). The final working concentration for each RNA was 0.1 μg/μL and the osmolarity of the pipette solution was 280-300 mOsm.

#### Single-cell electroporation in L2/3 of cortex under two-photon guidance

Anaesthetized mice with a half circular cortical window implanted over the target region of cortex were head-fixed under a 16X water immersion objective (NA = 0.8, 3 mm working distance, Nikon). A conductive, physiological gel (GEL G008 (FIAB), mixed with sucrose (179 mM) and NaCl (26.2 mM) for a final osmolarity of 280-300 mOsm), was added to the cranial window and an Ag/Cl ground electrode fixed inside immersive gel. Single cortical cells were electroporated as described previously^20^, with some important differences. Neurons were visualized by shadow imaging^23^ using a Femtonics FemtoSMART Resonant microscope connected to a Spectra-Physics InSight X3 two-photon laser tuned to 980 nm. Borosilicate glass pipettes (O.D.: 1.5 mm, I.D.: 0.86 mm, with filament, Sutter Instruments) were pulled to 6-12 MOhm using a P-97 micropipette puller (Sutter Instrument), filled with electroporation solution, and positioned with a micromanipulator (Luigs and Neumann) under two-photon guidance. Air pressure on the pipette was regulated by a manometer and set to 100-300 mBar for penetration of the dura. Once Alexa dye could be visualized in the brain, the pressure was reduced to 50-80 mBar, and the pipette was advanced into close vicinity of a cortical cell body in layer 2/3 (defined as cortical depth of 100-250 μm). The pressure was then reduced to 8-12 mBar, while the pipette was moved to be just adjacent to the membrane of the target cell body. The resistance of the pipette tip was monitored, and as soon as contact to the cell membrane was made (signified by an increase of 10-20% of tip resistance), the electroporation protocol was started. The cell was electroporated using a pulse train (1 s duration, 100 Hz, 0.5-ms pulse width) generated from an Axoporator 800B (Molecular Devices). As this device is no longer in production and unavailable, we performed a portion of our experiments using the NPI Electroporator with a connected PulsePal stimulation pulse train generator (Sanworks). The voltage amplitude of the pulse train was manually adjusted for each pipette, depending on the pipette resistance (Fig. S1). In this way, we ensured that the transmembrane current was never above 1.0 μA. The first measure of success of the electroporation was the spreading of dye (Alexa 594) into the targeted cell. After this was observed, the pipette was retracted carefully about 50-70 μm. Pressure was again increased to 50-80 mBar and, if the pipette tip was clear, more cells could be electroporated in the same fashion without changing the pipette. After the target cells were electroporated, the mouse was returned to a stereoscope for inspection and a new 4-mm round glass coverslip was affixed over the brain using cyanoacrylate glue for subsequent chronic two-photon imaging.

### Two-photon imaging in awake head-fixed mice

For two-photon calcium imaging as well as chronic monitoring of reporter expression, acclimatized and awake mice with cortical windows were head fixed under the two-photon microscope and allowed to run on a running wheel. Two-photon images were captured using the Femtonics MESc software and acquired using a 16X water immersion objective (NA = 0.8, 3 mm working distance, Nikon), with the microscope in resonant scanning mode and the laser tuned to 980 nm. For long-term tdTomato or GFP reporter tracking, exposure was fixed on the first day of recording by measuring laser power output as well as setting the photomultiplier tube detectors (Femtonics, high sensitivity GaAsP detectors) to a fixed gain value. This was maintained throughout the imaging series on the subsequent sessions. Image z-stacks were acquired averaging 80 times for each imaging plane spaced 5 μm apart.

### Whole-cell recordings in awake head-fixed mice

For awake whole-cell recordings, mice were used that had undergone acclimatization to being head-fixed in near-darkness on a running wheel. Additionally, a small cardboard shield was suspended above the body of the animal to prevent the tail from touching any of the recording equipment. As described above for single-cell electroporation, a mouse with a half-circular window over the cortical cells of interest was head-fixed under the microscope. The mouse was allowed to run on a running wheel while a diverse set of visual stimuli (see visual stimulus presentation below) was passively presented. A conductive physiological gel (GEL G008 (FIAB), mixed with sucrose (179 mM) and NaCl (26.2 mM) for a final osmolarity of 280-300 mOsm), was added to the cranial window and an Ag/Cl ground electrode was fixed inside immersive gel.

Neurons were visualized as described for single-cell electroporation and previously electroporated cells were targeted. In the case of target cells expressing tdTomato, their expression was used in conjunction with shadow imaging to direct the pipette to their location. Whole-cell patch pipettes (O.D.: 1.5 mm, I.D.: 0.86 mm, borosilicate glass with filament, Sutter Instrument) were pulled to 4-6 MOhm using a P-97 micropipette puller (Sutter Instrument) and filled with whole-cell internal solution: Voltage clamp, in mM: CsMeS [130], CsCl [4], NaCl [2], HEPES [10], EGTA [0.2], phosphocreatine-Tris [14], QX-314 [5], ATP-Mg [4], GTP-Tris [0.3], and Alexa Fluor 594 [0.025]; Current clamp, in mM: EGTA [0.2], K-gluconate [130], KCl [4], NaCl [2], HEPES [10], ATP-Mg [4], GTP-Tris [0.3], phosphocreatine-Tris [14] and Alexa Fluor 594 [0.025]. Intracellular solutions were calibrated to pH 7.25 and an osmolarity of 290 mOsm. Air pressure on the pipette was regulated by a manometer and set to 100-300 mBar for penetration of the dura. Once Alexa dye could be visualized in the brain, the pressure was reduced to 50-80 mBar, the pipette was advanced close to the target neuron, and the pressure was reduced to 30 mBar. Upon visual contact with the membrane, the positive pressure was released and a GΩ seal was formed. Slow and fast pipette capacitances were compensated, and whole-cell access was gained by applying a negative pulse of pressure. Recordings were done using a Multiclamp 700B amplifier digitized at 20 kHz. Access resistance was compensated as much as possible and monitored at the beginning of each recording and with each change of recording configuration. A-14-mV liquid junction potential was corrected in all voltage-clamp command signals. Cells were clamped either at-68 mV (to measure excitation) or 0 mV (to measure inhibition). Acceptable recordings were defined as having access resistances less than 60 MΩ.

### Visual stimulus presentation

During head-fixation for awake experiments (with the exception of current injection experiments), we displayed visual stimuli sequences for the mouse to activate visual cortical circuits. The right eye of the mouse was facing the center of an LCD screen (W: 47 cm, H: 27 cm) at a distance of 14.5 cm. Stimuli were generated, displayed and timed on a dedicated computer using custom written Python code. For awake whole-cell recordings, the stimuli consisted of a variety of grating textures, flashed on the screen (stimulus size = 60 degrees) for 2 s. The stimulus set contained vertical, horizontal, hyperbolic, spiral and polar gratings displayed randomly at different orientations and spatial frequencies.

## Data analysis

All data analysis was performed in custom-written routines in MATLAB (R2023b, Mathworks).

### Electroporation success rate

The electroporation success rate was determined by calculating the ratio of fluorescently labeled neurons on the day following the procedure to the total number of electroporated neurons. The success rate was initially computed for each individual field-of-view (FOV) or animal, and the resulting summary statistic across FOVs/animals was then presented. For electroporation of Cre mRNA, the success rate was determined on day 2 after electroporation.

### Quantification of fluorescence in electroporated neurons

To quantify fluorescence in neurons over time, a region of interest (ROI) encompassing each electroporated neuron was first manually selected in ImageJ^54^ on the imaging volume from the day following electroporation. An average projection of the three central Z-planes containing each neuron was then computed for every recording day. Raw fluorescence was extracted for each day and neuron by applying the original ROI to the projection image. To correct for baseline fluorescence, an additional ROI was selected in a region close to the electroporated cells on each imaging day, and this baseline fluorescence was subtracted from the neuron ROI fluorescence. Missing data points from days when imaging did not occur were linearly interpolated between previous and subsequent imaging sessions. If a cell was not identifiable on a given recording day, its corrected fluorescence was set to 0. To minimize day-to-day variability, fluorescence traces of each neuron were filtered with a Gaussian filter (standard deviation of two days) and normalized to their maximum fluorescence across days. For display purposes, the averaged trace was then again max normalized. For chronic imaging of Cre-induced tdTomato fluorescence, ROIs were initially selected on day 5. Curves were averaged across cells. For panels relating to neuronal visibility after electroporation, a neuron was considered visible if it had a fluorescence value of at least 5% of its maximum fluorescence. Half-lives of fluorescence decay were computed by fitting a simple exponential without offset to the fluorescence trace of each neuron independently. For double-labeling experiments, fluorescence in both the green and red channels was extracted for each neuron as described in the previous paragraph. Only the day after electroporation was considered and no normalization was applied. A neuron was considered double labeled if the ratio in fluorescence between the green and red channel was located between 0.1 and 10.

### CRISPR/Cas9 editing success rate

To assess the success rate of CRISPR/Cas9 editing of the *EGFP* gene, fluorescence traces were extracted and filtered from electroporated neurons as previously described. A similar number of non-electroporated control neurons in the local vicinity were also randomly selected and analyzed within each FOV. The fluorescence trace of each neuron was then normalized by day 1 fluorescence. To minimize the impact of global changes in imaging quality over time, all traces were additionally normalized by the mean fluorescence trace of the selected control cells in the same FOV. Successful *EGFP* knockouts were defined as those with a normalized fluorescence of less than 20% at the end of the experiment (day 30). In experiments utilizing two distinct guides against *EGFP*, imaging was limited to a few selected days. In these cases, the measured fluorescence at day 30 was normalized by the average fluorescence recorded on days 1 and 2 post electroporation. Knock-out success rate was initially computed for each individual FOV or animal, and the resulting summary statistic across FOVs/animals was then presented.

### Synaptic current event identification during voltage-clamp recordings

In order to identify fast synaptic inhibitory and excitatory currents in whole-cell voltage clamp recordings from awake head-fixed mice, raw traces were downsampled to 5 kHz and split into 7-s trials for both conditions. A template matching strategy was then employed, utilizing two distinct manually curated template events (one for inhibitory and one for excitatory event detection) that were cross correlated with the recording traces to identify putative events. The traces were preprocessed with a short (2 ms) Savitzky-Golay low-pass filter of polynomial order 2, followed by subtraction of a long median high-pass filter (inhibition: 70 ms, excitation: 200 ms) to correct for baseline shifts before template matching. Events with a template correlation of 0.5 or higher were considered for further analysis. The peak of the event was defined as the time point with maximum current within a 2-ms window, and the amplitude was calculated as the difference between the peak current and the minimum current within 8 ms before the peak but after the preceding peak. The peak timing and amplitude were recorded if they were above a minimum threshold (inhibition: 30 pA, excitation: 15 pA) to remove noise. The total charge was calculated as the sum of the current amplitudes per trial, and the event frequency was calculated as the average number of events per trial, irrespective of amplitude.

### Analysis of spontaneous and visually evoked activity during current clamp recordings

In awake current clamp recordings, each cell was continuously measured at 20 kHz for a total of 90 s. Action potential spikes were detected by identifying the peak of membrane potential fluctuations that surpassed at least the-20 mV threshold. The spike rate was defined as the total number of spikes with the 90-s period.

To determine depolarized and hyperpolarized membrane potential states, action potentials were first removed from voltage traces. This was achieved by determining the duration of each spike (from the points where the membrane potential reached 10% of its maximum amplitude in a period of 2.5 ms before the spike to 10 ms after the spike) and subsequently removing these periods from the voltage trace. Then, for each cell, the upper and lower baseline potentials were determined by taking respectively the mean of the membrane potentials above the 90% percentile and below the 10% percentile. The hyperpolarized state threshold was then defined as the sum of the lower baseline potential and a quarter of the difference between the upper and lower baseline potentials. Similarly, the depolarized state threshold was defined as the sum of the lower baseline potential and the half of the difference between the upper and lower baseline potentials. All timepoints corresponding to membrane potentials exceeding the depolarized state threshold were classified as depolarized states, while all timepoints corresponding to membrane potentials below the hyperpolarized state threshold were classified as hyperpolarized states.

### Analysis of intrinsic membrane properties during controlled current injections

In controlled current injections in awake animals, recordings were first downsampled to 20 kHz. For each current injection level, spikes were detected within the first 500 ms after onset of the current injection. Action potential spikes were detected by identifying the peak of membrane potential fluctuations that surpassed at least the +5 mV threshold. The spike rate was defined as the total number of spikes with the 500-ms period. Rheobase was defined as the lowest current injection level at which at least one spike was evoked within the 500-ms period.

To determine the input resistance, the baseline membrane potential was first computed as the median membrane potential in the 100-ms period before current injection. The median membrane potential during the current injection was then computed as steady state membrane potential for each current level. The first 60 ms after the start of the current injection were ignored to allow for the membrane potential to stabilize. For each current injection level, the input resistance was computed as the ratio between the difference of steady state and baseline membrane potentials and the input current. The final input resistance per cell was defined as the median input resistance for all recorded current injections levels ranging from-400 pA to-150 pA.

### Statistics

All statistical tests were performed in MATLAB (R2023b, Mathworks). The Wilcoxon rank-sum test using the exact method to compute the *p*-value was used. For repeated-measure designs, Kruskal–Wallis one-way analysis of variance followed by Dunn’s multiple comparison test were used. In all plots, the median and the interquartile range were used for display purposes, unless specified otherwise. All statistical tests were two-tailed and described in the figure legends. The value of *n* and what type of data it represented were specified for each figure in the corresponding figure legend. Significance levels were indicated by stars as follows: ns: *p* ≥ 0.05, ∗: *p* < 0.05, ∗∗: *p* < 0.01, ∗∗∗: *p* < 10^−3^.

