## Supplementary Information for "In vivo single-cell gene editing using RNA electroporation reveals sequential adaptation of cortical neurons to excitatory-inhibitory imbalance"

### Supplementary material

Figure S1: Details of electroporation parameters and intracellular dye persistence

Figure S2: Dynamics of EGFP expression in L2/3 neurons electroporated with EGFP mRNA

Figure S3: Single-cell mRNA electroporation success rates and off-target labeling

Figure S4: Reporter fluorescence timelines after knockout of EGFP via single-cell RNA electroporation

Figure S5: Voltage clamp recordings of control and GABA-A-R CRISPR electroporated neurons

Figure S6: Excitatory input, membrane and spiking properties of early-stage GABA-A-R CRISPR neurons

Video S1: Time-lapse recording of sequential single-cell RNA electroporation in layer 2/3 mouse cortical neurons.

Left panel displays GCaMP6s fluorescence signals, while right panel shows Alexa 594 dye delivery from the electroporation electrode. Seven individual neurons were electroporated in sequence, with elapsed time indicated in the bottom right corner.

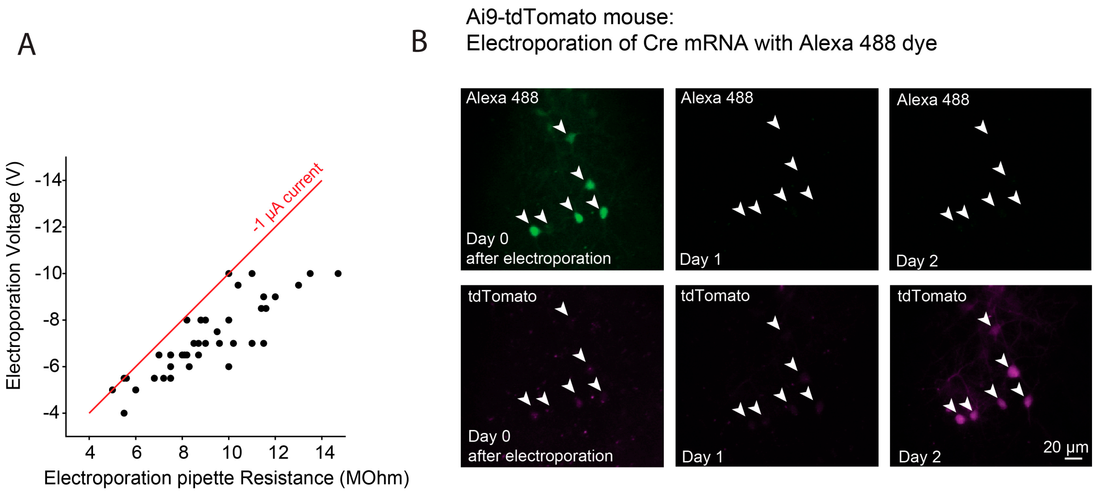


**Figure S1: Details of electroporation parameters and intracellular dye persistence**

**A:** Electroporation voltage plotted versus electroporation pipette resistance for each electroporation pipette used (*n* = 44 pipettes). The red line depicts a final electroporation current at the tip of the electrode of -1 µA. **B:** Example images of neurons in an Ai9-tdTomato mouse (which expresses tdTomato in the presence of Cre) electroporated with Cre mRNA and Alexa 488 dye, recorded on the day of electroporation (left: day 0) and the two following days. (Top) Green images show Alexa 488 dye fluorescence and (bottom) magenta images show tdTomato expression.

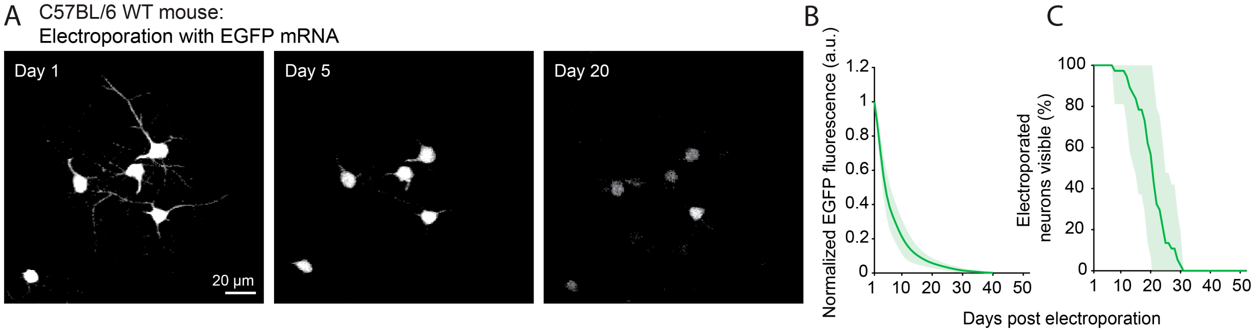


**Figure S2: Dynamics of EGFP expression in L2/3 neurons electroporated with EGFP mRNA**

**A:** Expression of EGFP in L2/3 neurons after single-cell electroporation of EGFP mRNA, imaged over multiple weeks. **B:** Mean EGFP fluorescence intensity (green line) of electroporated neurons plotted against days since electroporation of reporter mRNA. $n$ = 37 neurons from four imaging sites in three mice. Green shading, STD across neurons. **C:** Fraction of visible EGFP mRNA electroporated neurons plotted against days since electroporation. Green shading, STD across neurons. Same cells as in **B**.

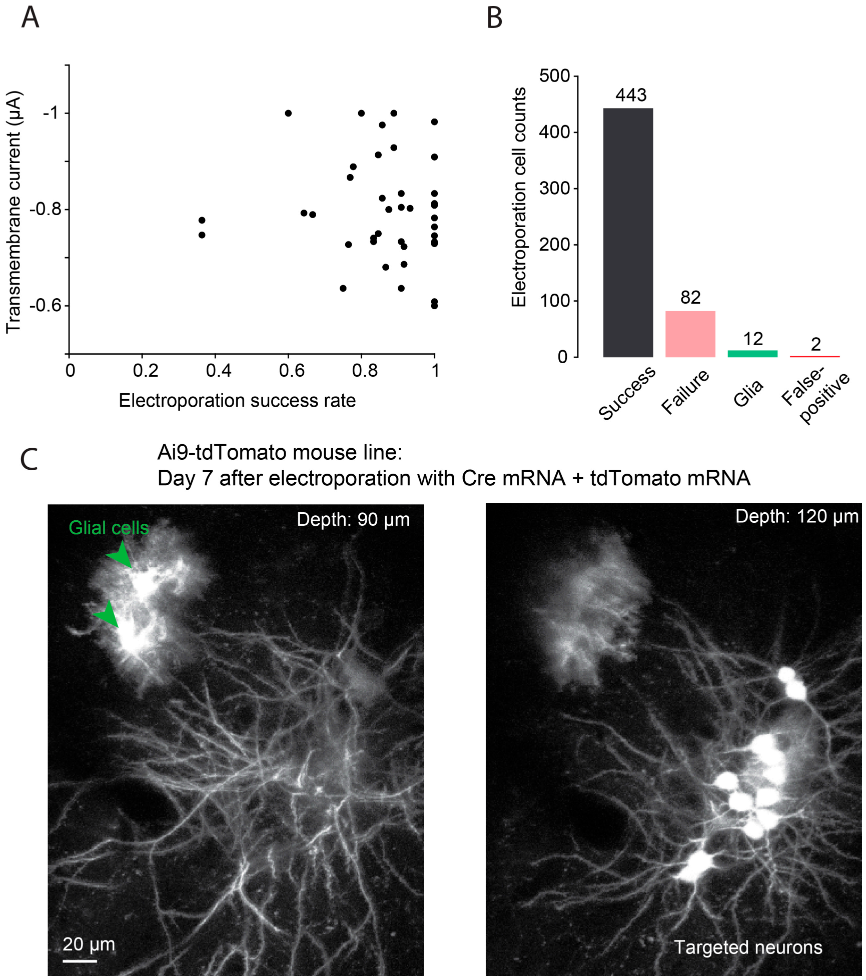


**Figure S3: Single-cell mRNA electroporation success rates and off-target labeling**

**A:** Transmembrane current during electroporation (calculated from electroporation voltage and pipette resistance) plotted against electroporation success rate. Each data point corresponds to an electroporation pipette (*n* = 44 pipettes). **B:** Outcomes of all electroporation attempts (*n* = 525 electroporation attempts). Success was defined as an electroporation that resulted in the targeted neuron expressing either the mRNA reporter or Cre-dependent genomic reporter. Glial cell false-positive labeling was identified through visual inspection (see lower panels in **C**). False-positive neurons were identified when the number of labeled neurons exceeded the number of electroporations performed. **C:** Example images after electroporation of L2/3 neurons in an Ai9-tdTomato mouse with Cre mRNA. Different planes of the same region are shown to depict the visible morphological differences between untargeted glial cells expressing tdTomato after electroporation (left: green arrows) compared to L2/3 neurons that were targeted via the electroporation pipette (right).

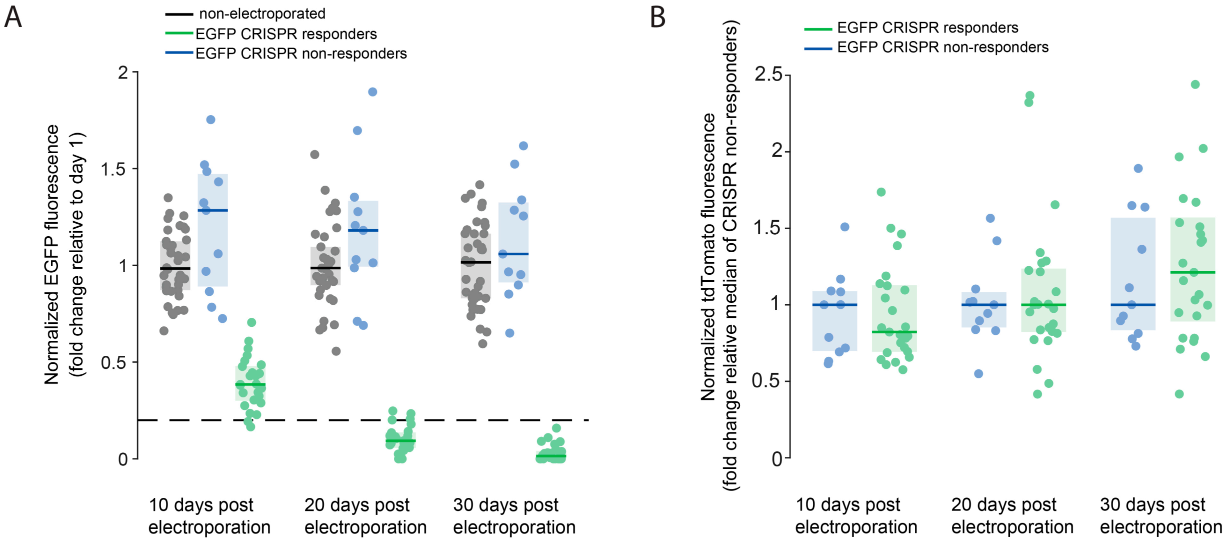


**Figure S4: Reporter fluorescence timelines after knockout of *EGFP* via single-cell RNA electroporation**

**A:** EGFP fluorescence of EGFP$^+$ L1 cortical neurons after the electroporation of tdTomato reporter mRNA, Cas9 mRNA and gRNA against *EGFP* in GAD67-*EGFP* mice plotted at different days post electroporation. Fluorescence values normalized to day 1 fluorescence. CRISPR editing is considered successful (green) if day 30 fluorescence intensity is less than 20% (black dashed line) of day 1 fluorescence intensity, while CRISPR editing is considered unsuccessful (blue) otherwise. $n$ = 36 cells from five imaging sites in four mice. Grey data points depict non-electroporated EGFP$^+$ L1 cortical neurons in the same animals ($n$ = 38 cells from five imaging sites in four mice). Each data point corresponds to a cell. Black, blue and green solid lines, median; gray, blue and green shadings, IQR across neurons. **B:** Reporter tdTomato fluorescence of the same electroporated EGFP$^+$ L1 cortical neurons shown in **A** plotted at different days post electroporation. Green and blue data points correspond to a cell each and depict successful and unsuccessful CRISPR action against *EGFP*, respectively. Fluorescence values normalized to the median tdTomato fluorescence of CRISPR non-responders on the corresponding day. Blue and green solid lines, median; blue and green shadings, IQR across neurons. One point (EGFP CRISPR responder 30 days post electroporation, value 4.151) was omitted from the plot for display reasons.

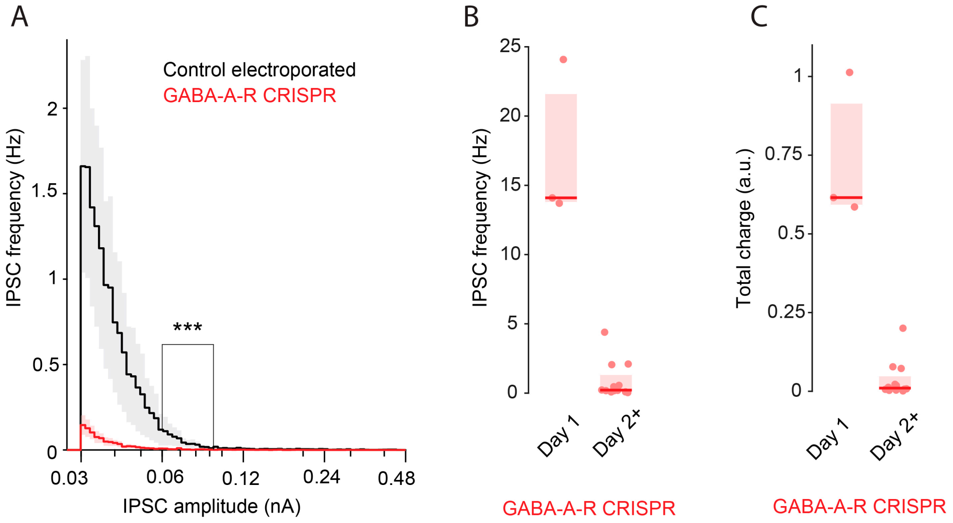


**Figure S5: Voltage clamp recordings of control and GABA-A-R CRISPR electroporated neurons**

**A:** Mean population distributions of IPSC strength in control electroporated neurons (black, $n = 5$ neurons in three mice) and GABA-A-R CRISPR electroporated neurons (red, $n = 12$ neurons in seven mice, recorded 2 days or more after electroporation). Gray and red shadings, SEM across neurons. $p = 7.90 \times 10^{-9}$, two-sample Kolmogorov-Smirnov test. **B:** IPSC frequency recorded in GABA-A-R CRISPR neurons at different timepoints after electroporation (at day 1: $n = 3$ neurons in two mice; at day 2+: $n = 12$ neurons in seven mice). Each data point corresponds to a cell. Red lines, median; red shadings, IQR across neurons. **C:** IPSC charge recorded in GABA-A-R CRISPR neurons at different timepoints after electroporation (at day 1: $n = 3$ neurons in two mice; at day 2+: $n = 12$ neurons in seven mice). Each data point corresponds to a cell. Red lines, median; red shadings, IQR across neurons.

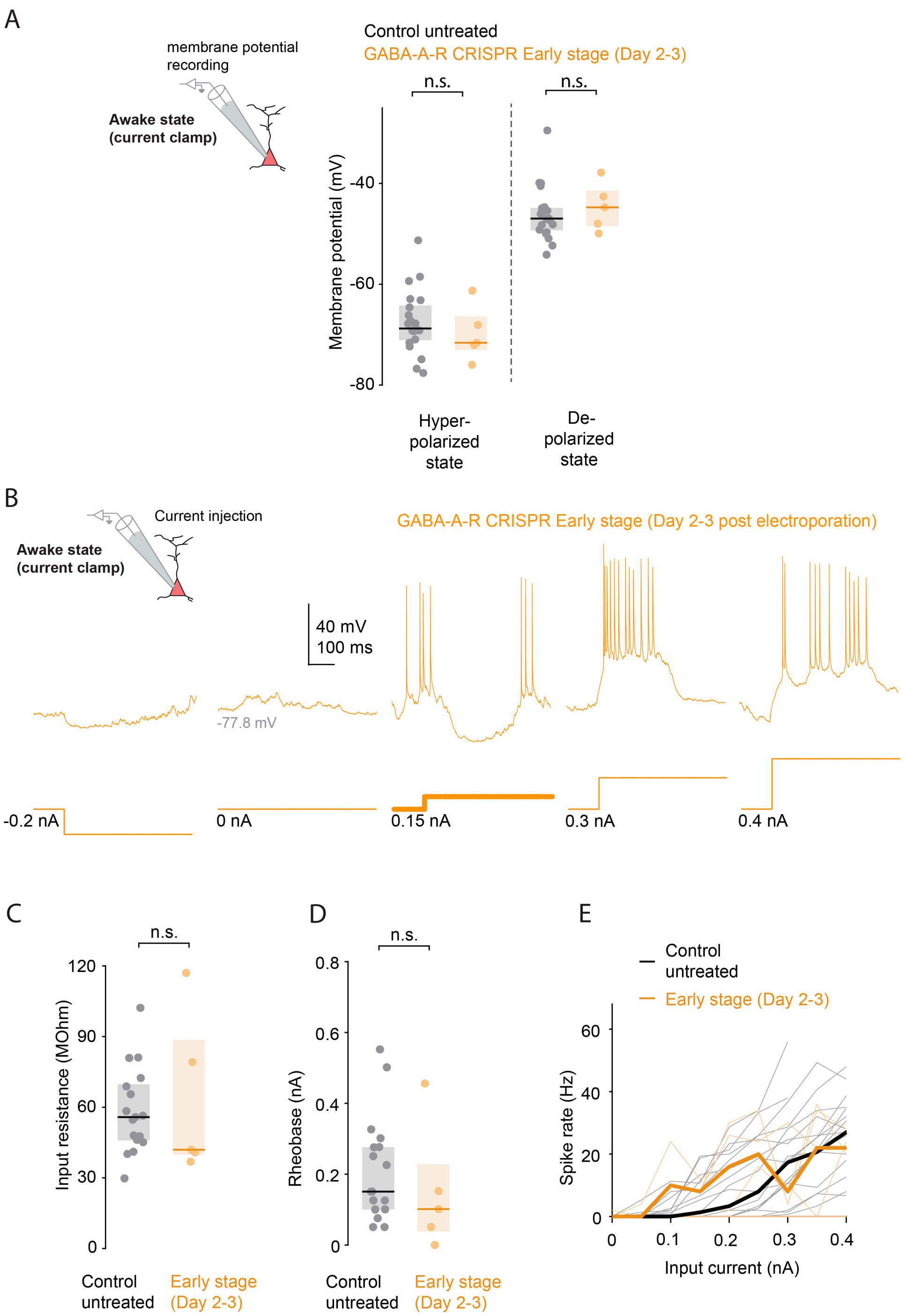


**Figure S6: Excitatory input, membrane and spiking properties of early-stage GABA-A-R CRISPR neurons**

**A:** Hyperpolarized (left) and depolarized (right) membrane potential of untreated neurons (black, $n$ = 21 neurons in nine mice) and early-stage GABA-A-R CRISPR neurons (yellow, $n$ = 5 neurons in two mice). Each data point corresponds to a cell. Black and yellow lines, median; gray and yellow shadings, IQR across neurons. Hyperpolarized membrane potential: $p = 0.410$, depolarized membrane potential: $p = 0.447$, Wilcoxon rank-sum test. **B:** Awake whole-cell current clamp recording of membrane potential in an early-stage GABA-A-R CRISPR L2/3 neuron (2-3 days after electroporation) during current injections. The corresponding current step trace is displayed below the membrane potential trace. Bold current trace indicates rheobase. **C:** Input resistance of untreated neurons (black, $n = 19$ neurons in nine mice) and early-stage GABA-A-R CRISPR neurons (yellow, $n = 5$ neurons in two mice). Each data point corresponds to a cell. Black and yellow lines, median; gray and yellow shadings, IQR across neurons. $p = 0.704$, Wilcoxon rank-sum test. **D:** Rheobase of untreated neurons (black, $n = 19$ neurons in nine mice) and early-stage GABA-A-R CRISPR neurons (yellow, $n = 5$ neurons in two mice). Each data point corresponds to a cell. Black and yellow lines, median; gray and yellow shadings, IQR across neurons. $p = 0.436$, Wilcoxon rank-sum test. **E:** Median spike rate (bold line) of untreated neurons (black, $n = 19$ neurons in nine mice) and early-stage GABA-A-R CRISPR neurons (yellow, $n = 5$ neurons in two mice) in response to current injections. Each pale line corresponds to a cell.
